# Microbial dysbiosis and its implications for disease in a genetically depauperate species

**DOI:** 10.1101/653220

**Authors:** Alexandra L. DeCandia, Julie L. King, Bridgett M. vonHoldt

## Abstract

THE host-associated microbiome is increasingly recognized as a critical player in health and immunity. When commensal microbial communities are disrupted, dysbiosis can contribute to disease pathogenesis and severity. Santa Catalina Island foxes (*Urocyon littoralis catalinae*) present an ideal case study for examining dysbiosis in wildlife due to their depauperate genomic structure and extremely high prevalence of ear canal tumors. Although the precise cause is yet unknown, infection with ear mites (*Otodectes cynotis*) has been linked to chronic inflammation, which is associated with abnormal cell growth and tumor development. Given the paucity of genomic variation in these foxes, other dimensions of molecular diversity, such as commensal microbes, may be critical to host response and adaptation. We therefore characterized the host-associated microbiome across six body sites of Santa Catalina Island foxes, and performed differential abundance testing between healthy and mite-infected ear canals. We found that mite infection was significantly associated with reduced microbial diversity and evenness, with the opportunistic pathogen *Staphylococcus pseudintermedius* dominating the ear canal community. These results suggest that secondary bacterial infection may contribute to the sustained inflammation associated with tumor development. Uncovering high abundance of *S. pseudintermedius* provides critical insight into the pathogenesis of this complex system, as the emergence of antibiotic resistant strains remains a concern of the medical, veterinary, and conservation communities. Through use of culture-independent sequencing techniques, this study contributes to the broader effort of applying a more inclusive understanding of molecular diversity to questions within wildlife disease ecology.

## INTRODUCTION

Channel Island foxes (*Urocyon littoralis*) are among the most genetically depauperate mammals in the world. Monomorphic at the majority of loci, island foxes exhibit extremely low levels of genome-wide heterozygosity ranging from 0.14% on San Nicolas to 4.08% on Santa Catalina (Aguilar et al., 2004; Robinson et al., 2016). Although these foxes show little evidence of inbreeding depression, disease remains a salient threat to long-term persistence, particularly on Santa Catalina Island (SCA), which hosts a population of resident humans and domestic animals year-round. For example, an outbreak of canine distemper virus (CDV) in 1999 led to a large-scale mortality event that caused a 95% population decline on the East End of the island (King, Duncan, & Garcelon, 2014; Timm et al., 2009). In response, a conservation management program was immediately launched in 2000 that included translocation from the West End, captive breeding, and vaccination against CDV. This enabled SCA foxes (*U. l. catalinae*) to recover, with annual monitoring and vaccination underway to prevent future outbreaks.

Ceruminous gland tumors, including both carcinomas and adenomas, are also found at extremely high rates among SCA foxes. Between 2001 and 2008, 48.9% of dead foxes had tumors in their ear canals, with 52.2% of mature live foxes sampled in 2007-2008 showing signs of tumor growth (Vickers et al., 2015). Although the underlying mechanisms are yet unknown, the forerunning conceptual model links chronic inflammation to tumorigenesis. This model hypothesizes that the presence of ear mites (*Otodectes cynotis*) leads to otitis externa, followed by ceruminous gland hyperplasia (CGH) and tumor development (Vickers et al., 2015). The association between sustained inflammation and tumor growth has been observed in domestic dogs (*Canis familiaris*) and cats (*Felis catus*), where mite-infection and chronic otitis externa preceded aural tumor diagnosis (London et al., 1996; Sula, 2012). These connections were further substantiated with SCA foxes, where experimental acaricide treatment led to marked decreases in mite burden, inflammation, CGH, and mite-specific IgG antibodies (Moriarty et al., 2015).

Although the links between ear mites, inflammation, and tumorigenesis are becoming clearer, little is known about the role that the host-associated microbiome plays in this system. Commensal microbes are known to compete with pathogenic invaders, regulate immunity through metabolite production, and aid the resolution of immune responses through altered gene expression and immune signaling (Berbers, Nierkens, van Laar, Bogaert, & Leavis, 2017; Buffie & Pamer, 2013; K Honda & Littman, 2012; Kenya Honda & Littman, 2016; Naik et al., 2012). Dysbiosis, or disruption of these communities, has been linked to autoimmunity, inflammation, and disease. Examples include humans and dogs suffering from otitis externa and allergic skin disease, as well as pigs (*Sus scrofa domesticus*) experimentally infected with sarcoptic mange (*Sarcoptes scabiei* var. *suis*). In each case, infected individuals exhibited lower levels of microbial diversity, altered community abundance, and increased incidence of pathogenic *Staphylococcus spp.* (Bannoehr & Guardabassi, 2012; Bradley et al., 2016; Kong et al., 2012; Ngo, Taminiau, Fall, Daube, & Fontaine, 2018; Rodrigues Hoffmann et al., 2014; Swe, Zakrzewski, Kelly, Krause, & Fischer, 2014). This evidence suggests that disrupted microbes at the skin barrier allow for colonization of opportunistic, secondary bacterial infections that further stress the host immune system.

Culture-based methods have previously been used to characterize ear canal microbes in SCA foxes, but these efforts may have been biased by methodological constraints. When ear canal bacteria were cultured from foxes with and without ceruminous gland tumors, no associations were found between pathogenic bacterial strains and tumor presence (Vickers et al. 2015). However, culture-based methods are known to underrepresent microbial taxa in canine ear canals and other body sites (Ngo et al., 2018; Rodrigues Hoffmann et al., 2014). Further, by focusing on tumor presence, the study design may have excluded meaningful patterns of dysbiosis that occur during earlier stages of the conceptual model linking ear mites, inflammation, CGH, and tumorigenesis.

In the present study, we used next generation sequencing to characterize microbial communities associated with ear mite infection. Using culture-independent methods, we focused on the first stage of the conceptual model put forth by Vickers et al. (2015). We hypothesized that the presence of ear mites would be associated with secondary bacterial infection and decreased microbial diversity that contribute to the chronic inflammation associated with CGH and tumor development in this system. Sampling five additional epidermal sites, we characterized microbial communities across haired and mucosal body surfaces to establish a healthy baseline of microbes present on this subspecies of endemic foxes.

## MATERIALS AND METHODS

### Study area

Santa Catalina Island has an area of 194km^2^ and is located 32km southwest of Long Beach, California (33° 24’ N, 118° 24’ W). Approximately 88% of the island is owned and managed by the Catalina Island Conservancy, with the remaining 12% owned and managed by the Santa Catalina Island Company (11%) and the City of Avalon (1%). Nearly 4,000 human residents live on SCA year-round, and approximately 800,000 people visit the island each year. Catalina is mountainous, with a Mediterranean climate characterized by hot, dry summers and mild winters. Major plant communities include coastal sage scrub, oak woodland, island chaparral, and grasslands (Schoenherr, Feldmeth, & Emerson, 1999).

### Sample and data collection

We captured and collected biological samples from SCA foxes during two consecutive field seasons under the authorization of a U.S. Fish and Wildlife Service (USFWS) Federal 10(a)1(A) permit (No. TE 090990-2), a Memorandum of Understanding with the California Department of Fish and Wildlife (CDFW), and Scientific Collecting Permits (No. 005821 and No. 009858) issued by CDFW. Our procedures were reviewed and approved by the Princeton University Institutional Animal Care and Use Committee (Princeton IACUC #2046A-18).

In 2017 and 2018, we opportunistically collected samples from wild foxes trapped during annual monitoring efforts led by the Catalina Island Conservancy. Foxes were sampled if they were greater than six months old and had not been treated with Ivermectin during the previous 10 months. Welded-wire box traps (Model #106 Tomahawk Live Trap Company) were set and baited with a combination of cat kibble, canned cat food, and loganberry lure. Traps were placed along roadside transects at 250–300m intervals with 32-49 sites used daily. We checked all traps each morning for four consecutive days before moving them to a new location.

Foxes were muzzled and manually restrained while we collected samples and baseline data, including passive integrated transponder tag number, sex, age, weight, body condition, and ear canal infection status. Foxes were assigned to one of five age groups (0-4) using tooth wear patterns as described by Wood (1958); however, the age of most animals was known from previous capture history (range: 0-10 years). All foxes were examined, sampled, and released at the site of capture.

We observed ear canals otoscopically and assigned ear mite severity scores ranging from 0 to 4 as defined by Vickers et al. (2015) and Moriarty et al. (2015). The presence and severity of ear canal wax, *liquor puris*, pigmentation, inflammation, and ceruminous gland tumors (or nodules) were also assessed and recorded. After the initial assessment, samples were collected using a sterile BBL™ Culture Swab™ inserted into the ear canal and rotated approximately 180 degrees 10 times before being returned to its pre-labeled applicator. When necessary, additional non-sterile cotton swabs were used to remove residual keratin debris post-sampling, which allowed for a more detailed otoscopic examination to detect any visible abnormalities. All ear canals were then treated topically with the acaricide Ivermectin (IVOMEC^®^ Injection, 0.05 ml of 1% solution, Merial Ltd.). Following every examination, otoscope specula were disinfected with 70% isopropyl alcohol and stored submerged in 2% chlorhexidine diacetate solution (Nolvasan® Solution, Zoetis Inc.).

For the remaining body sites sampled, we rubbed the tip of a sterile BBL™ swab on the skin (or as close to the skin as possible) 40 times, rotating the swab tip by a quarter every 10 times before placing the swab back into its pre-labeled sterile applicator. We kept sample swabs in an insulated container with ice packs until storing them at −80°C at the end of each day. All traps were scrubbed and cleaned weekly using a 2% chlorhexidine diacetate solution to prevent disease and parasite transmission when traps were moved to a new location.

Between two field seasons, we sampled 76 unique individuals, with 17 foxes sampled both years. During November and December 2017, we swabbed 30 foxes at six body sites that included ear canal, external ear, nostril, lip commissure, axilla, and perianal area (Supplemental Figure S1). We also sampled 24 foxes at two body sites that included ear canal and external ear, for a total of 228 samples collected in the first field season. During October through December 2018, we swabbed four foxes at all six body sites (*N.B.,* one fox had an extra ear canal sample taken) and 35 foxes at both left and right ear canals, for a total of 95 samples. After field collection, all samples were stored at −80°C until DNA extraction.

### Microbial DNA extraction and amplicon sequencing

We extracted DNA following a modified DNeasy PowerSoil Kit (Qiagen Inc.) protocol. We used sterile scissors to isolate the swab tips directly into the PowerBead tubes, which were then disrupted for two cycles on a Qiagen TissueLyser II: 1) 12 minutes at 20 shakes/second and 2) an additional 12 minutes at 20 shakes/second after we added 60µL of C1 solution. We followed the standard manufacturer protocol for subsequent steps, with a modified elution for 10-15 minutes in 60µL C6 buffer pre-heated to 70°C. To ensure no contamination, we used a sterile swab tip as an extraction blank during each set of extractions. We concentrated extracts to 20µL in a Vacufuge and quantified double-stranded DNA with a high-sensitivity Qubit™ fluorometer. We standardized high concentration samples to 2.5ng/µL and included samples with concentrations as low as 0.110ng/µL.

We used 96 unique combinations of barcoded forward (n=8) and reverse (n=12) primers that target the 16S ribosomal RNA (rRNA) V4 region to amplify and tag each DNA sample (Caporaso et al., 2011). PCR reactions included 5µL HiFi HotStart ReadyMix (KAPA Biosystems), 3.2µL primer mix (1.25µM), and 1.8µL template DNA and followed cycling conditions consisting of: initial denaturation of 94^°^C/3 min followed by touchdown cycling for 30 cycles of [94°C/45s, 80-50°C/60s, 72°C/90s] decreasing 1°C each cycle, followed by 12 cycles of [94°C/45s, 50°C/60s, 72°C/90s], finishing with a final extension of 72°C/10min. We used Agencourt AMPure XP magnetic beads to remove nonspecific binding in the PCR products and eluted in a final volume of 15µL low TE (10mM Tris-HCl pH 7.5, 0.1 mM EDTA). We quantified PCR product using Quant-iT™ PicoGreen™ dsDNA assays and visualized 3µL of each sample on a 1% agarose gel to confirm amplification of the target region. We pooled 96 uniquely barcoded libraries and completed a selection for fragments between 300-400nt in size using Agencourt AMPure XP magnetic beads. We collected paired-end amplicon sequence data (2×150nt) on an Illumina MiSeq in the Princeton University Genomics Core Facility.

### Bioinformatics and data processing

We demultiplexed raw sequencing reads and allowed for a single nucleotide mismatch between the observed and expected sequence tags using a barcode splitter for paired-end, dual-indexed data in the online platform *Galaxy* (Afgan et al., 2018). We then imported demultiplexed reads into *QIIME 2* v2019.1 (Caporaso et al. 2010; https://qiime2.org) for filtering with the *dada2 denoise-paired* plugin (Callahan et al., 2016). This plug-in implements a model-based approach to correct probable sequencing errors, remove chimeric sequences, trim 13 low quality bases at the beginning of each read (flags *--p-trim-left-f* and *--p-trim-left-r*), and merge paired-end reads. Denoising is an alternative to operational taxonomic unit (OTU) construction and enables finer-scale resolution of microbial variation than traditional cluster-based methods, such as *de-novo, closed reference*, and *open reference* clustering implemented with the *qiime vsearch* function (Rognes et al. 2016).

### Alpha and beta diversity

In order to calculate phylogenetic diversity metrics, we constructed a rooted phylogenetic tree with the *alignment* and *phylogeny* functions in *QIIME 2*. We aligned all representative sequences, masked highly variable positions, and created a rooted phylogenetic tree using *fasttree,* with the root placed at the midpoint between the longest branches. We then estimated alpha and beta diversity metrics for samples rarefied to the maximum sampling depth that included all samples (depth=6381 sequences). We employed the following functions for diversity estimations: *core-metrics-phylogenetic, alpha, alpha-rarefaction, alpha-group-significance*, and *beta-group-significance*. Alpha (within-sample) diversity metrics included: the Chao 1 index to represent species richness; Shannon’s diversity index for species abundance and evenness; and Pielou’s evenness metric for species equitability. We used the Kruskal-Wallis test to assess statistical significance of within-group diversity. To examine beta (between-sample) diversity, we used the Bray-Curtis dissimilarity index to measure species abundance and estimated two phylogeny-based UniFrac distances (unweighted distances for species presence; weighted distances for species presence and abundance). We visualized differences through principal coordinate analysis (PCoA) implemented with the *EMPeror* plugin (Vázquez-Baeza, Pirrung, Gonzalez, & Knight, 2013). We assessed statistical significance of between-group differences using a multivariate analysis of variance with permutation (*PERMANOVA*; Anderson 2001).

### Taxonomic analysis

To examine the taxonomic composition of each sample, we trained a Naïve Bayes classifier on reference sequences from the Greengenes 13_8 database clustered at 99% similarity and trimmed to the region targeted by our primers (Bokulich et al., 2018; DeSantis et al., 2006). We then used the *classify-sklearn* function in the *QIIME 2 feature-classifier* plugin to assign taxonomic labels to our representative sequences with the newly trained classifier (Bokulich et al., 2018). In addition to standard visualizations, we used the Bray-Curtis and Euclidean distance metrics to hierarchically cluster samples with the “average” linkage method (Hunter, 2007).

### Differential abundance testing

Given the statistical constraints of working with compositional data, we used two methods explicitly designed for differential abundance testing: analysis of composition of microbes (ANCOM) and gneiss balances. ANCOM calculates pairwise log-ratios between taxa to detect features that differ in abundance between groups, with the underlying assumption that only a few features change (Mandal et al., 2015). Gneiss calculates balances (or log transformed ratios) at each internal node of a tree relating all microbial features to one another (Morton et al., 2017). This enables identification of subcommunities that differ in abundance, which are tested further with linear regression models (see below).

For these analyses, we applied additional filters to our feature table to only include features with a minimum total frequency of 50 and minimum number of samples of 10. We then created a composition artifact, which contains feature frequencies per sample, and ran ANCOM for ear canals grouped by mite infection status (binary infected versus uninfected). We performed this analysis at both the class and feature levels, and queried results using NCBI BLASTn (Altschul, Gish, Miller, Myers, & Lipman, 1990).

For gneiss, we used correlation clustering to create a tree hierarchy that related all microbial taxa to one another, and visualized the tree using the Interactive Tree of Life (iTOL) v3 online tool (Letunic & Bork, 2016). We input the composition artifact into *gneiss ilr-hierarchical* function to calculate balances at each internal node with the isometric log ratio (ILR) transform. We used an ordinary least squares (OLS) regression on these balances, with fixed effects specified as the sampling area, sex, age class, weight, body condition, and ear mite presence. We additionally constructed a linear mixed effects (LME) model to account for resampled individuals and included fox ID as a random effect. We used *gneiss balance-taxonomy* to identify which taxa were present in balances showing differential abundance between infection groups, in order to identify the feature(s) driving the observed differences.

## RESULTS

### Amplicon sequencing and data processing

We sequenced 252 samples (external ear, n=33; ear canal, n=127; lip commissure, n=33; nostril, n=16; axilla, n=9; perianal area, n=34) that represented 76 foxes (samples per fox ranged from 1-10, with median=3 and mode=2). We obtained 11,493,554 raw sequences, of which we retained 8,354,655 sequences after denoising (Supplemental Table S1). This dataset contained 21,698 unique taxonomic features. In comparison, OTU-based methods yielded 9,709 features with *de novo* clustering, 6,946 features with closed-reference clustering, and 12,296 features with open-reference clustering, with percentage identity thresholds of 97%. We removed 18 samples from the denoised dataset due to erroneous treatment with Ivermectin (n=13), sampling near (rather than inside) the ear canal (n=1), unknown ear mite burden (n=2), and laboratory processing error (n=2). We created filtered feature tables from this pruned dataset for downstream statistical analyses. The “body” feature table (total n=186; external ear, n=32; ear canal, n=69; lip commissure, n=30; nostril, n=15; axilla, n=9; perianal area, n=31) included samples from all six body sites, but excluded ear canals from foxes with at least one mite-infected ear. The “ear canal” feature table (n=119) included only ear canal swabs, regardless of infection status.

### Skin microbiome of Santa Catalina Island foxes

#### Alpha and beta diversity significantly differ between body sites

We observed significant differences in alpha (Kruskal-Wallis test; Chao 1 index, *H*=69.852, *p*<0.001) and beta (PERMANOVA; Bray-Curtis dissimilarity index, *pseudo-F*=5.140, *p*=0.001) diversity between body sites. With sequencing depth rarefied to 6381, species richness was highest at haired sites (external ear=394.63±151.46 and axilla=365.15±127.40), with decreasing richness observed at the ear canal (327.92±173.98), nostril (270.36±147.14), perianal area (162.00±59.03), and lip commissure (149.31±75.30; Figure 1A). We observed a similar pattern using the Shannon index (*H*= 23.242, *p*<0.001), with nostril swabs exhibiting the lowest diversity (4.386±1.542), perhaps due to low species evenness (nostril=0.558±0.154; *H*=8.800, *p*=0.117; Supplemental Figures S2, S3). Principal coordinate plots of beta diversity (Figure 1B, Supplemental Figure S4) showed lip commissure and perianal samples as two distinct clusters, with all other body sites (axilla, ear canal, external ear, and nostril) overlapping one another.

**Figure 1.**
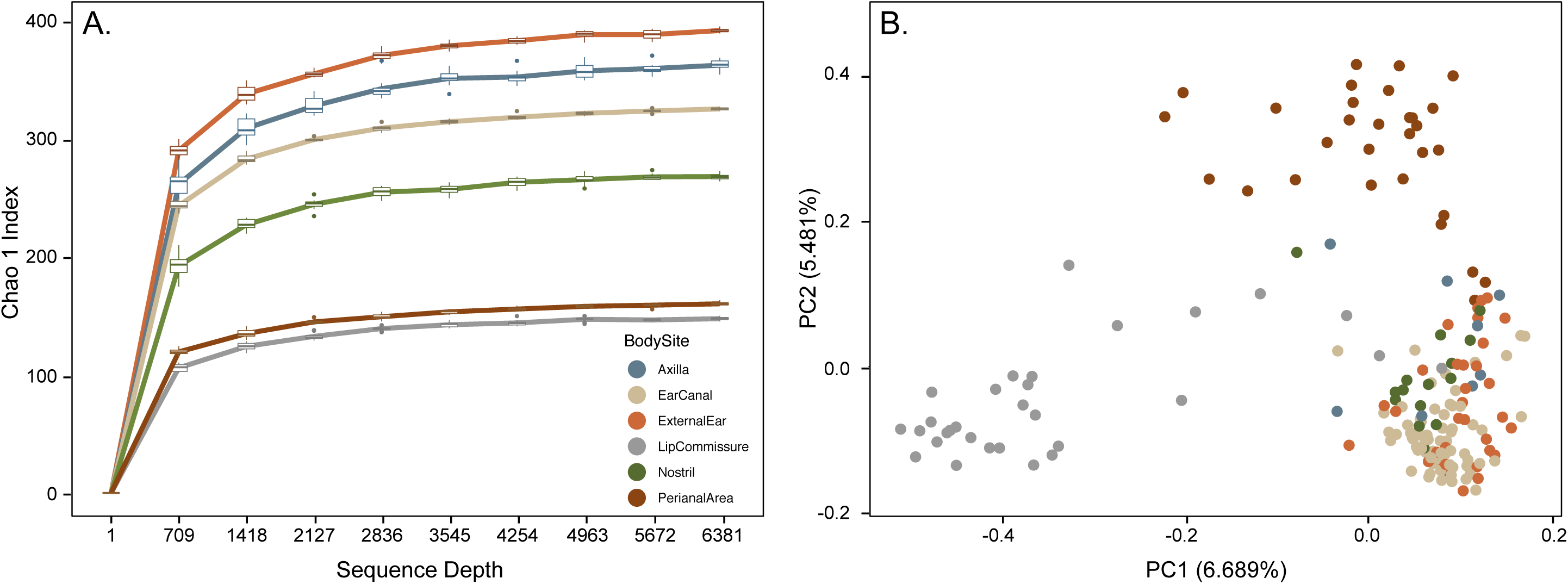
Alpha (Kruskal-Wallis test; Chao 1 Index, *H*=69.852, *p*<0.001) and beta (*PERMANOVA*; Bray-Curtis dissimilarity index, *pseudo-F*=5.140, *p*=0.001) diversity metrics significantly differed by body site. (A) Rarefaction curve of the Chao 1 Index, where lines represent mean values across samples and iterations (n=10), and box plots represent mean values across samples for each iteration. (B) Principal coordinate analysis (PCoA) plot of the Bray-Curtis dissimilarity index.

#### Relative abundance underlies differences in taxonomic composition between sites

The four body sites included in the overlapping region of the PCoA plot (Figure 1B) showed similar taxonomic composition and abundance (Figure 2, Supplemental Figure S5). Firmicutes was the most abundant phylum detected (axilla=37.33%, nostril=29.35%, external ear=27.05%, and ear canal=20.00%) with Proteobacteria (range=14.93-23.24%), Bacteriodetes (range=10.13-16.66%), and Actinobacteria (range=8.50-17.73%) generally present in even abundances at those sites. In contrast, Proteobacteria dominated the lip commissure community (44.53%), whereas Bacteriodetes was the most abundant taxon at the perianal area (34.92%). Following these patterns, hierarchical clustering of body sites using both Bray-Curtis and Euclidean distance metrics placed lip and perianal area in their own grouping, with (ear canal + external ear) more closely related to the (nostril + axilla) grouping (Supplemental Figure S6).

**Figure 2.**
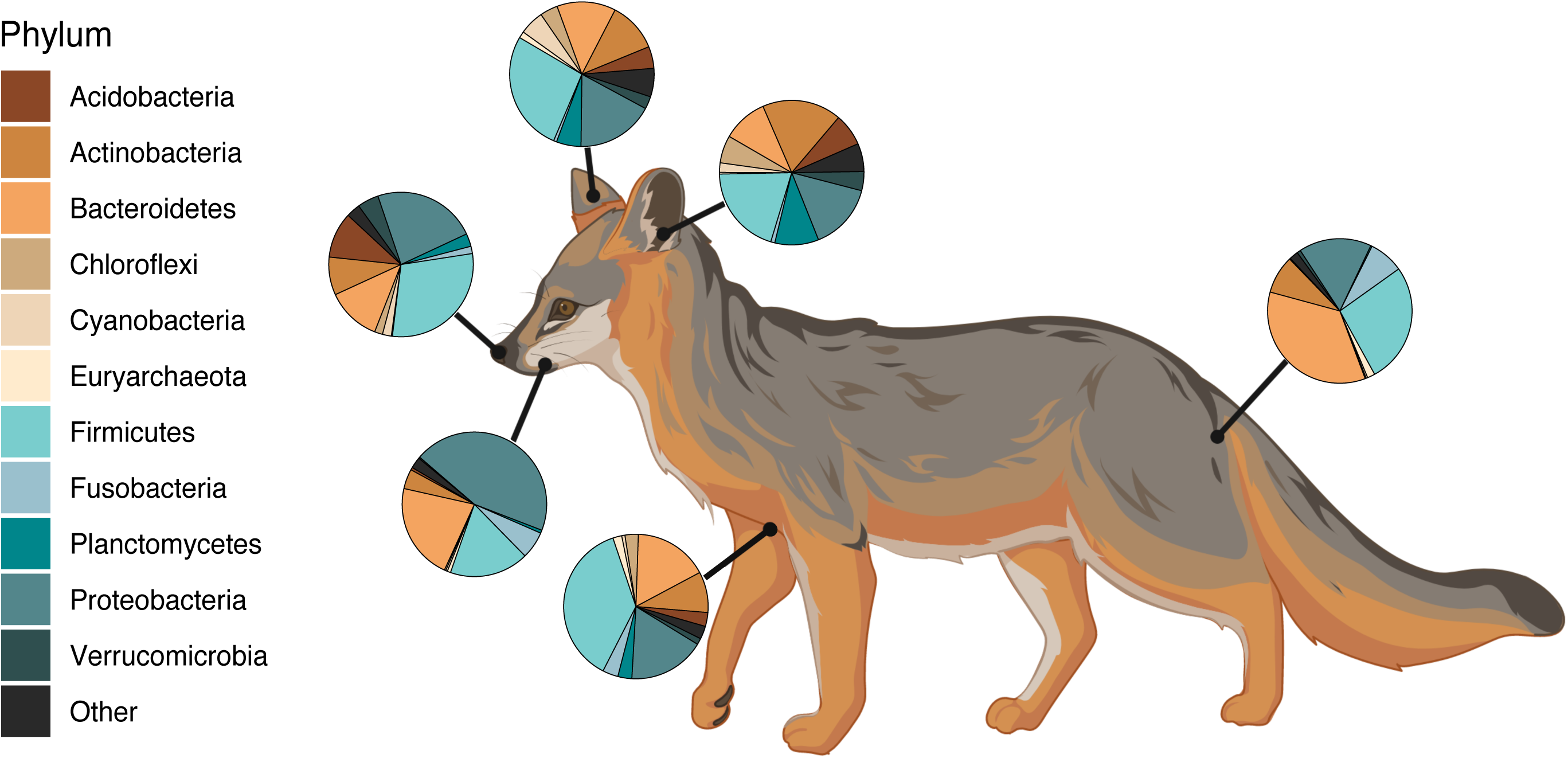
Taxonomic composition at each body site at the phylum level. Body sites include: ear canal, external ear, nostril, lip commissure, axilla, and perianal area. Illustration created with BioRender (https://biorender.com/).

### Ear canal diversity of mite-infected and uninfected foxes

#### Significant correlations detected between ear canal phenotypes

We performed pairwise Chi-square tests using base functions in *R* to assess the independence of ear canal phenotypes as measured by the binary presence of mites, wax, *liquor puris*, ceruminous gland tumors (nodules), pigment, and inflammation (Supplemental Table S2). All phenotypes were significantly correlated (*p*-values below the FDR-corrected threshold 0.015; Benjamini & Yekutieli 2001), with the exception of nodules and mites (*X*^*2*^=1.597, *p*=0.206), wax (*X*^*2*^=1.319, *p*=0.251), and *liquor puris* (*X*^*2*^=4.856, *p*=0.028). Given that nodules were heavily skewed towards the absence class and the remaining12/15 tests returned non-independence between variables, we used mite presence as our phenotype of interest for downstream analyses.

#### Mite-infected ears exhibit decreased species richness and evenness

Alpha diversity was significantly reduced in mite-infected ear canals. This trend was consistent across metrics, with species richness (Kruskal-Wallis; Chao1 Index, *H*=25.635, *p*<0.001; Figure 3A, Supplemental Figures S7A, B), abundance and evenness (Shannon Index, *H*=31.481, *p*<0.001; Supplemental Figures S7C,D), and equitability (Pielou’s Evenness Metric, *H*=27.848, *p*<0.001; Figure 3B, Supplemental Figures S7E,F) all significantly lower. Beta diversity similarly differed by mite infection status (*PERMANOVA*; Bray-Curtis dissimilarity index, *pseudo-F*=8.139, *p*=0.001; unweighted UniFrac distance, *pseudo-F*=4.281, *p*=0.001; and weighted UniFrac distance, *pseudo-F*=8.942, *p*=0.001; Figure 3C, Supplemental Figure S8). Samples loosely clustered by infection status in the Bray-Curtis PCoA plot (Figure 3C), where PC1 explained 16.810% of the variation and was significantly associated with mite presence (one-way ANOVA, *F*= 59.210, *p*<0.001).

**Figure 3.**
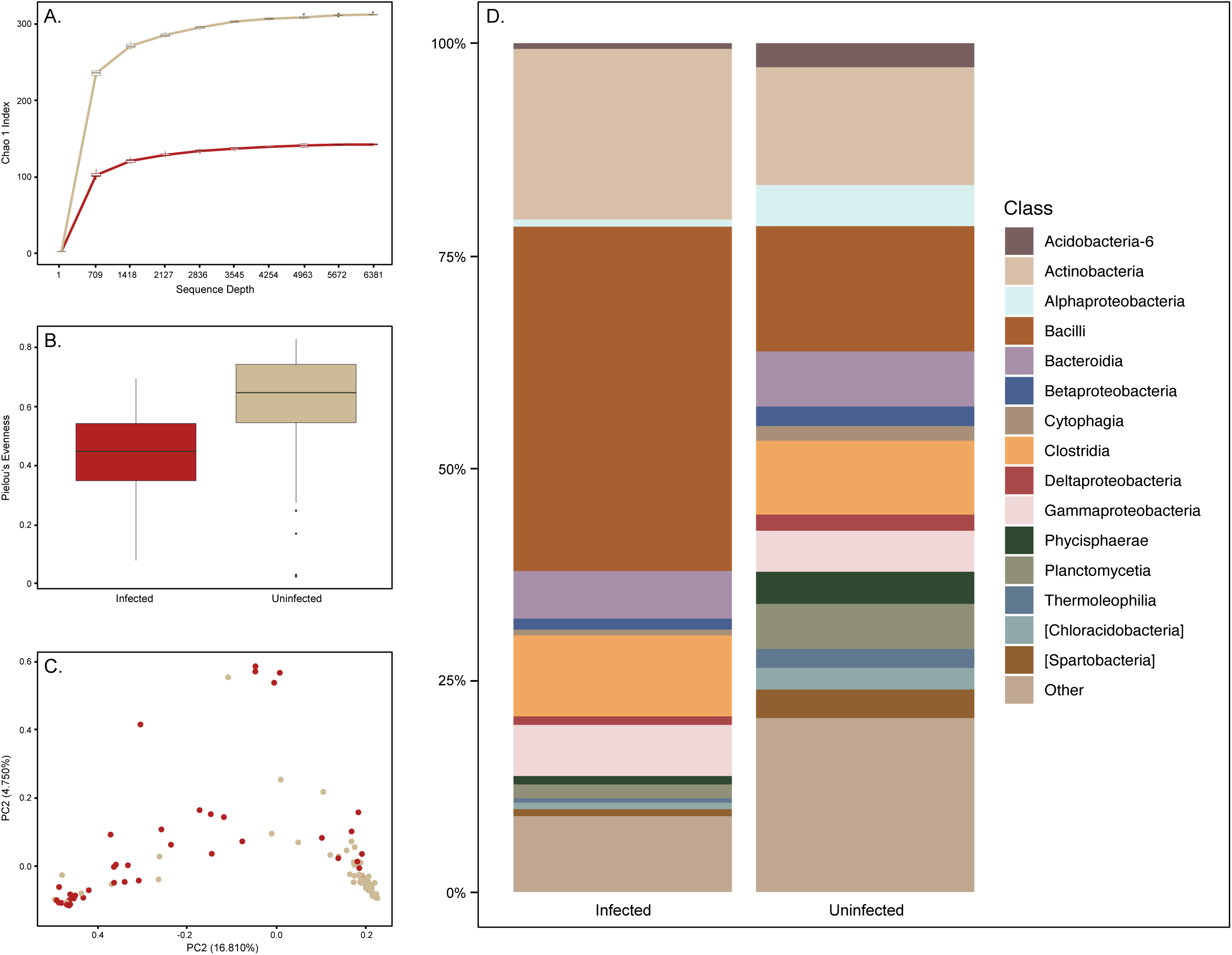
Microbial communities sampled in mite-infected and uninfected ear canals significantly differed across alpha diversity metrics measuring (A) species richness (Kruskal-Wallis Test; Chao 1 Index, *H*=25.635, *p*<0.001) and (B) equitability (Pielou’s Evenness Metric, *H*=27.848, *p*<0.001), and beta diversity metrics measuring (C) abundance (*PERMANOVA*; Bray-Curtis dissimilarity index, *pseudo-F*=8.139, *p*=0.001). (D) Taxonomic composition exhibited the same classes of microbes in both infection groups, with Bacilli far more abundant in mite-infected ear canals.

#### Taxonomic composition skewed towards Bacilli in mite-infected ear canals

Taxonomic composition and abundance differed with mite presence (Figure 3D). Uninfected ear canals had a higher number of taxa present at relatively even abundances (range 1.73-14.75%, all other taxa summed to 20.51%), with Bacilli (14.75%) and Actinobacteria (13.87%) the two dominant classes. In mite-infected ears, these two classes were disproportionately abundant (Bacilli=40.56% and Actinobacteria=20.09%), with all others present in far lower abundance (range 0.51-9.55%, all other taxa summed to 8.96%). Results obtained using our full (n=119, 1-3 samples per fox) and subsampled (n=73, 1 sample per fox) datasets were nearly identical (Supporting Information). We therefore used the full dataset for downstream analyses to maximize sample inclusion, being sure to note any discordant results.

#### Staphylococcus pseudintermedius (class Bacilli) drives differential abundance between groups

Bacilli was the only taxon with a significantly different abundance between infected and uninfected ear canals (*W*=98) using analysis of composition at the class level. When we performed this analysis at the feature level, feature 3f0449c545626dd14b585e9c7b2d16f4 (*W*= 418) was shown to drive this pattern. This feature had high sequence similarity using NCBI BLASTn to *Staphylococcus pseudintermedius* (Supplemental Table S3).

In the OLS regression model constructed with gneiss balances (*r*^*2*^=0.092), ear mite presence was the variable of largest effect (*r*^*2*^ *diff=*0.023; Supplemental Table S4). We identified one balance that was statistically significant with respect to ear mite infection (balance y01, *p*<0.001). This balance displayed a higher proportion of Bacilli in mite-infected ear canals (Figure 4A), and contained feature 3f0449c545626dd14b585e9c7b2d16f4, previously identified as *S. pseudintermedius*, as the primary taxon differing in proportion between mite-infected (mean=0.471) and uninfected (mean=0.120) foxes. Correlation-clustering grouped this feature with two others in the tree hierarchy (Figure 4B): feature 4cdbebb204548db0a0f4d1860eab7fef, which matched to *Corynebacterium spp.* using NCBI BLASTn (Altschul et al., 1990), and feature 13e52764e1c08c4fd9902769bf9e022e, which matched to *Enterococcus spp.* Both features were present in higher proportions in infected (*Corynebacterium* mean=0.058, *Enterococcus* mean= 0.019) versus uninfected (*Corynebacterium* mean=0.009, *Enterococcus* mean= 0.014) ear canals.

**Figure 4.**
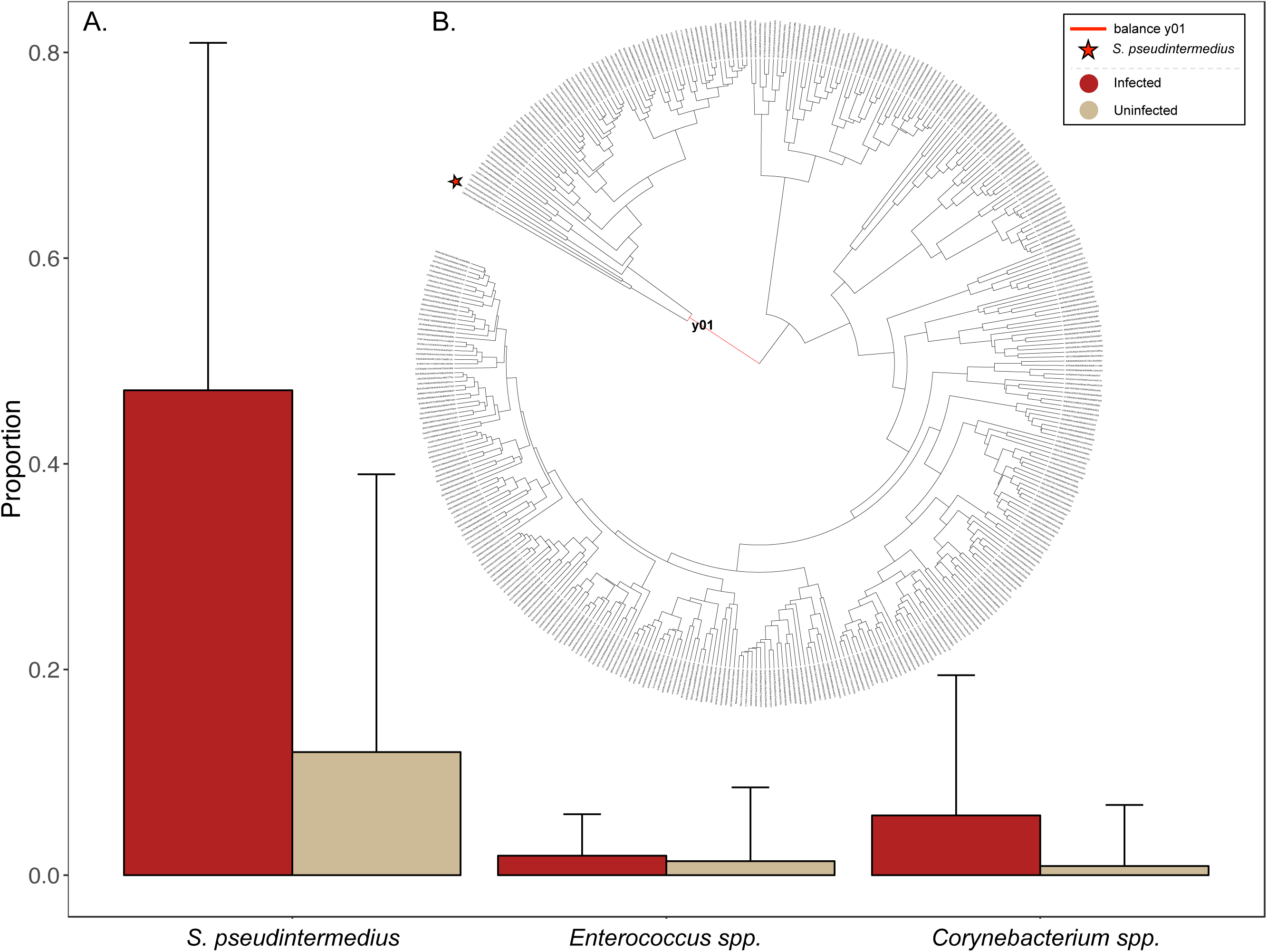
(A) Proportion bar plots for three taxonomic features contained in both gneiss balances associated with ear mite infection. *S. pseudintermedius* had the largest difference in proportion between mite-infected and uninfected ear canals, with *Enterococcus sp.* and *Corynebacterium sp*. clustered with *S. pseudintermedius* in the (B) hierarchy relating all microbial taxa to one another. Balance y01 is highlighted in red, and *S. pseudintermedius* is indicated with a star.

To account for the 36 foxes sampled multiple times, we constructed a LME model with fox ID specified as a random effect. We found that the same balance significantly differed in abundance (balance y01, *p<0.001*), with a higher proportion of *S. pseudintermedius* in infected ear canals. The small number of samples per fox (range 1-3) resulted in difficulty fitting this model, as this analysis is optimized for datasets with at least 3 repeated measures per individual. We therefore constructed a second OLS model with a subsampled dataset that included only one sample per fox selected at random (Supporting Information). In this analysis, the balance significantly associated with mite infection returned the same result: *S. pseudintermedius* (Bacilli) had significantly higher relative abundance (mean=0.627) in mite-infected ear canals compared to uninfected ear canals (mean=0.146). In the subsampled tree hierarchy, only *Enterococcus spp.* (Bacilli) was clustered with *S. pseudintermedius.*

## DISCUSSION

In the absence of genetic diversity, other dimensions of molecular variation can provide critical insights into population health, disease pathogenesis, and, ultimately, the evolution of host-parasite systems. As such, we used culture-independent, next generation sequencing to characterize the host-associated microbiome in an endemic population of Channel Island foxes. Focusing on six body sites, we established a baseline of healthy microbial communities across both haired and mucosal surfaces. We additionally reported patterns of dysbiosis in ear canals infected with ear mites, as measured by decreased microbial diversity and increased incidence of *S. pseudintermedius.* This suggests a possible link between secondary bacterial infection and the chronic inflammation associated with CGH and ceruminous gland tumors in SCA foxes.

### Skin microbiome of SCA foxes

As the importance of host-associated microbiomes is increasingly recognized (McKenney, Koelle, Dunn, & Yoder, 2018; Parfrey, Moreau, & Russell, 2018), there has been a surge in studies characterizing microbial communities in diverse sets of taxa. Within marine and terrestrial mammals, recent examples include Humpback whales (Apprill et al., 2014), Tasmanian devils (Cheng et al., 2015), lowland gorillas (Gomez et al., 2015), bats (Avena et al., 2016; Ingala, Becker, Bak Holm, Kristiansen, & Simmons, 2019), orcas (Hooper et al., 2019), and Antarctic fur seals (Grosser et al., 2019), among others. Phenotypes of interest range from behavior to disease to ecology; yet, microbial natural history, or the characterization of taxonomic composition and diversity of the host-associated microbiome, remains a key objective of the field.

We contributed to this effort by sequencing microbial communities at six body sites of endemic SCA foxes. We found significant differences in alpha and beta diversity across body sites, with haired skin exhibiting higher species richness than mucosal surfaces and mucocutaneous junctions, as seen in domestic dogs (Hoffmann et al., 2014). Dominant phyla included Firmicutes, Proteobacteria, Bacteriodetes, and Actinobacteria across most sites, with additional phyla abundant in nostril (Acidobacteria) and lip/perianal (Fusobacteria) areas. These phyla are commonly reported for domestic (Hoffmann et al., 2014), terrestrial (Avena et al., 2016; Wu et al., 2017), and marine (Grosser et al., 2019) mammal systems, although relative abundances differ by host species and body site. These represent the core microbiota of SCA foxes, and may serve as a baseline for future efforts monitoring population health and community composition following disturbance, such as disease.

### Ear canal diversity of mite-infected and uninfected foxes

Mite-infected ear canals exhibited patterns of diversity indicative of microbial dysbiosis. Alpha diversity metrics measuring species richness, abundance, equitability, and evenness were all significantly reduced in mite-infected ear canals. Similar reductions have been reported for dogs suffering from atopic dermatitis (Bradley et al., 2016; Hoffmann et al., 2014), with changes in beta diversity evident in dogs with otitis externa (Ngo et al., 2018). We observed similar differences in beta diversity using the Bray-Curtis dissimilarity index, which suggested that species abundance may contribute to the differences observed between infection groups.

Taxonomic composition confirmed skewed abundance between groups, with Bacilli the dominant class associated with mite infection. Using multiple approaches for differential abundance testing, we discovered that *S. pseudintermedius* was the bacterial species consistently and significantly more abundant in mite-infected ear canals. This result mirrors numerous systems, including inflammatory conditions affecting human and canine skin and ears (Bradley et al., 2016; Hoffmann et al., 2014; Kong et al., 2012; Lappan et al., 2018; Ngo et al., 2018; Williams & Gallo, 2015). Increased prevalence of *Staphylococcus spp.* has also been reported for pigs experimentally infected with *S. scabiei* var. *suis* mites (Swe et al., 2014), where the authors documented changes in skin microbial conditions before, during, and after experimental mite infection. In the present study, *S. pseudintermedius* commonly co-occurred with *Enterococcus spp.* and *Corynebacterium spp.* However, the relative abundances of these additional taxa differed by a smaller magnitude, with *Enterococcus spp.* the only co-occurring taxon in the subsampled analysis. We therefore concluded that *S. pseudintermedius* was the primary taxon driving the observed differences between mite-infected and uninfected ear canals.

*S. pseudintermedius* is a common commensal microbe primarily found on species within the family Canidae, which includes foxes and domestic dogs (Bannoehr, Franco, Iurescia, Battisti, & Fitzgerald, 2009). Considered an opportunistic pathogen, *S. pseudintermedius* can quickly proliferate when the balance of microbial communities is disrupted by allergic skin disease, such as atopic dermatitis (Fazakerley et al., 2009), or an external stimulus, such as surgery (Bannoehr & Guardabassi, 2012). Second to the fungal pathogen *Malassezia pachydermatis, Staphylococcus spp.* are the leading bacterial cause of otitis externa in dogs, with *Enterococcus spp., Corynebacterium spp.*, and *Pseudomonas spp.* among others commonly cultured (Ngo et al., 2018). Often associated with ear, skin, and post-operative infections in dogs, cats, and occasionally humans, *S. pseudintermedius* forms biofilms that are remarkably difficult to eradicate after proliferation (Pompilio et al., 2015). As a result, this taxon is associated with chronic inflammation across host systems. The emergence of antibiotic resistant strains remains a prominent concern of the medical, veterinary, and conservation communities (Sasaki et al., 2007; Weese & van Duijkeren, 2010).

Within the context of SCA foxes, these results suggest an amendment to the conceptual model put forth by Vickers et al. (2015). We now hypothesize that ear mite infection *and* secondary bacterial infection with the opportunistic pathogen *S. pseudintermedius* both contribute to the sustained inflammation associated with CGH and tumor development on Santa Catalina Island.

Through use of culture-independent methods, we uncovered patterns of microbial dysbiosis associated with ear mite infection that are consistent with expectations drawn from similar disease systems (*e.g.,* Swe et al. 2014). Ongoing treatment with Ivermectin has already proven effective at decreasing mite burdens and inflammation (Moriarty et al., 2015), and will hopefully lead to lower tumor rates as the years progress. In the interim, these additional insights into the microbial pathogenesis of this complex host-parasite system enable consideration of treatment with pro- or antibiotics, if necessary (Lappan et al., 2018). Identification of *S. pseudintermedius* as the bacteria underlying these changes also enables monitoring for the emergence of antibiotic resistant strains (Weese & van Duijkeren, 2010). Particularly in an endemic island population lacking genetic diversity, an outbreak of antibiotic resistant bacteria can pose a serious conservation risk.

Broadly, this study contributes to the ongoing call for the application of microbial sequencing techniques within the fields of molecular and disease ecologies and wildlife conservation (DeCandia, Dobson, & vonHoldt, 2018; Hauffe & Barelli, 2019; Trevelline, Fontaine, Hartup, & Kohl, 2019). By sequencing the microbiome at six body sites of healthy and mite-infected foxes, we established a baseline of healthy microbiota while also identifying patterns of dysbiosis associated with their perturbation. Applying similar techniques to wild and captive systems around the world will not only further our understanding of the role of host-associated microbiomes in health and disease, but may also provide insights useful for population monitoring, management, and conservation. Particularly in species lacking genetic diversity, other dimensions of molecular variation, such as commensal microbial communities, can contribute to the health, viability, and evolutionary potential of depauperate populations. These implications render the study of host-associated microbiomes an important and growing area within the field of molecular ecology.

## ACKNOWLEDGEMENTS

We would like to thank Calvin Duncan, Lara Brenner, Rebekah Rudy, and Emily Hamblen from the Catalina Island Conservancy for aiding sample collection, and Yuanyuan Cheng, Elizabeth Grice, and Aline Rodrigues Hoffman for their advice on swabbing protocols at the outset of this study. We would additionally like to thank Lindy McBride for use of her Qiagen TissueLyser II for DNA extraction, and Wei Wang and Jessica Wiggins from the Princeton University Genomics Core Facility for their assistance optimizing the library preparation protocol and initial stages of data processing. Funding for this study was provided by the Theodore Roosevelt Memorial Fund of the American Museum of Natural History and by the Catalina Island Conservancy. This material is based upon work supported by the National Science Foundation Graduate Research Fellowship under Grant No. DGE1656466.

## DATA ACCESSIBILITY

Upon acceptance, demultiplexed forward and reverse reads will be deposited to the NCBI Sequence Read Archive.

## AUTHOR CONTRIBUTIONS

ALD, JLK, and BMvH designed the study; JLK collected the samples; ALD conducted the laboratory work; ALD processed and analyzed the data; ALD, JLK, and BMvH prepared, contributed to, and approved the final manuscript.

