## Supplemental for "Microbial dysbiosis and its implications for disease in a genetically depauperate species"

**Supporting Information 1: Supplemental Tables and Figures**

**Supplemental Figure S1.** We sampled six body sites that included (1) ear canal, (2) external ear, (3) nostril, (4) lip commissure, (5) axilla, and (6) perianal area.

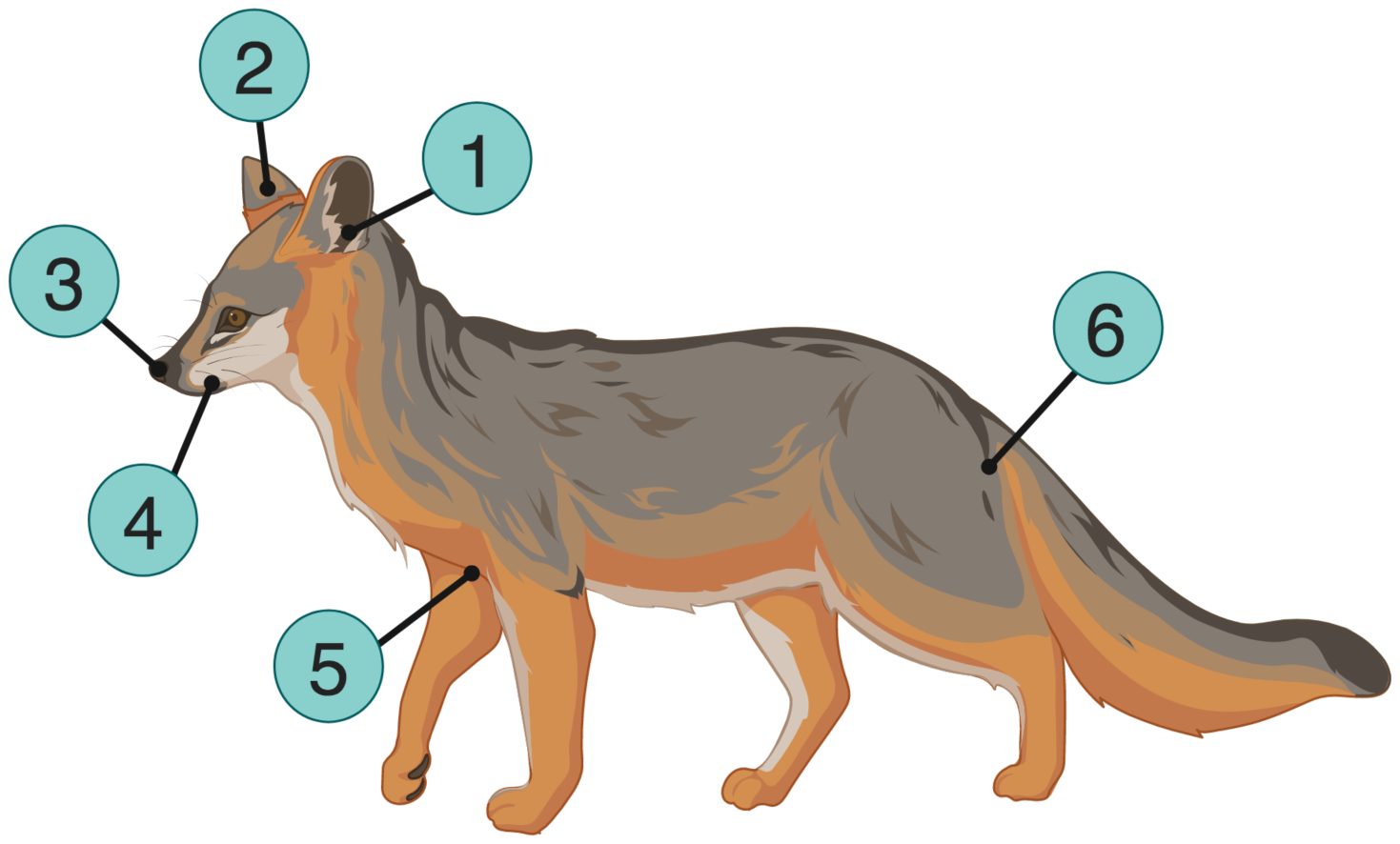

**Supplemental Table S1.** Denoising statistics by sample from raw sequences through quality filtering, dada2 denoising, paired-end merging, and chimera removal.

| **Sample** | **Raw** | **Filtered** | **Denoised** | **Merged** | **Non-chimeric** |
| --- | --- | --- | --- | --- | --- |
| Fox-1 | 35660 | 29180 | 29180 | 27314 | 27270 |
| Fox-2 | 48324 | 41560 | 41560 | 38309 | 38262 |
| Fox-3 | 32007 | 27981 | 27981 | 26906 | 26877 |
| Fox-4 | 57932 | 43618 | 43618 | 41580 | 41424 |
| Fox-5 | 40582 | 34253 | 34253 | 32404 | 32295 |
| Fox-6 | 59082 | 51059 | 51059 | 48538 | 48500 |
| Fox-7 | 70205 | 60503 | 60503 | 57709 | 57454 |
| Fox-8 | 39270 | 36907 | 36907 | 35679 | 35179 |
| Fox-9 | 67023 | 54172 | 54172 | 51734 | 51550 |
| Fox-10 | 45003 | 31230 | 31230 | 28392 | 28298 |
| Fox-11 | 34448 | 22202 | 22202 | 20830 | 20830 |
| Fox-12 | 42706 | 38028 | 38028 | 36944 | 36426 |
| Fox-13 | 42202 | 36164 | 36164 | 34855 | 34248 |
| Fox-14 | 33611 | 29007 | 29007 | 28524 | 28303 |
| Fox-15-1† | 39396 | 31685 | 31685 | 28927 | 28829 |
| Fox-15-2† | 43836 | 34197 | 34197 | 31113 | 31011 |
| Fox-16-1† | 48387 | 38371 | 38371 | 35029 | 35011 |
| Fox-16-2† | 40185 | 36249 | 36249 | 34204 | 34132 |
| Fox-17 | 34138 | 25230 | 25230 | 23009 | 23009 |
| Fox-18 | 80611 | 68860 | 68860 | 66052 | 65769 |
| Fox-19 | 38961 | 30276 | 30276 | 26545 | 26492 |
| Fox-20 | 41579 | 28546 | 28546 | 24523 | 24491 |
| Fox-21 | 49525 | 43220 | 43220 | 41864 | 41646 |
| Fox-22 | 67103 | 47964 | 47964 | 43618 | 43536 |
| Fox-23 | 68052 | 58684 | 58684 | 57136 | 56613 |
| Fox-24 | 47806 | 36031 | 36031 | 33714 | 33659 |
| Fox-25 | 36002 | 30375 | 30375 | 29215 | 27078 |
| Fox-26 | 90280 | 74307 | 74307 | 70567 | 70276 |
| Fox-27 | 45838 | 24427 | 24427 | 20363 | 20332 |
| Fox-28 | 39139 | 33713 | 33713 | 32819 | 32584 |
| Fox-29 | 44808 | 32873 | 32873 | 31571 | 31404 |
| Fox-30 | 31267 | 24473 | 24473 | 21893 | 21868 |
| Fox-31 | 40127 | 32096 | 32096 | 30479 | 30441 |
| Fox-32 | 51276 | 43085 | 43085 | 39925 | 39875 |
| Fox-33 | 46762 | 34149 | 34149 | 30120 | 30092 |
| Fox-34 | 45551 | 34743 | 34743 | 32500 | 32416 |
| Fox-35 | 47767 | 37807 | 37807 | 36263 | 35880 |
| Fox-36 | 48275 | 42440 | 42440 | 40082 | 40066 |
| Fox-37 | 44907 | 40482 | 40482 | 39058 | 38945 |
| Fox-38 | 54957 | 43720 | 43720 | 39509 | 39459 |
| Fox-39 | 53057 | 40770 | 40770 | 38665 | 38175 |
| Fox-40 | 38413 | 22563 | 22563 | 19530 | 19492 |
| Fox-41 | 32360 | 24793 | 24793 | 22654 | 22636 |
| Fox-42 | 62821 | 51683 | 51683 | 48994 | 48513 |
| Fox-43 | 27325 | 16964 | 16964 | 15084 | 15073 |
| Fox-44 | 46074 | 24064 | 24064 | 21683 | 21645 |
| Fox-45 | 41310 | 35357 | 35357 | 33398 | 33184 |
| Fox-46 | 61690 | 48758 | 48758 | 45761 | 45705 |
| Fox-47 | 42469 | 31082 | 31082 | 25739 | 25666 |
| Fox-48 | 52002 | 29729 | 29729 | 24451 | 24352 |
| Fox-49 | 40105 | 34198 | 34198 | 32820 | 32609 |
| Fox-50 | 62453 | 46830 | 46830 | 42556 | 42518 |
| Fox-51 | 42061 | 37169 | 37169 | 36206 | 35901 |
| Fox-52 | 52567 | 44137 | 44137 | 39908 | 39643 |
| Fox-53 | 51706 | 43770 | 43770 | 40695 | 40407 |
| Fox-54 | 63115 | 54278 | 54278 | 51797 | 51337 |
| Fox-55 | 38798 | 35451 | 35451 | 33515 | 33163 |
| Fox-56 | 53763 | 45903 | 45903 | 43476 | 43384 |
| Fox-57 | 39238 | 27494 | 27494 | 26052 | 26022 |
| Fox-58 | 45964 | 42498 | 42498 | 41314 | 41055 |
| Fox-59 | 36140 | 31766 | 31766 | 30593 | 30529 |
| Fox-60 | 57290 | 53584 | 53584 | 53203 | 53203 |
| Fox-61 | 57262 | 50386 | 50386 | 48784 | 48161 |
| Fox-62 | 52425 | 44427 | 44427 | 41945 | 41694 |
| Fox-63 | 74787 | 56924 | 56924 | 53443 | 53415 |
| Fox-64 | 43864 | 39200 | 39200 | 37048 | 36909 |
| Fox-65 | 43418 | 35266 | 35266 | 33355 | 33313 |
| Fox-66 | 73183 | 59252 | 59252 | 54983 | 53999 |
| Fox-67 | 53324 | 46867 | 46867 | 45914 | 45871 |
| Fox-68 | 42671 | 40176 | 40176 | 39237 | 38941 |
| Fox-69 | 37847 | 25634 | 25634 | 23757 | 23746 |
| Fox-70 | 37503 | 29283 | 29283 | 27820 | 27606 |
| Fox-71 | 52159 | 38117 | 38117 | 31482 | 31381 |
| Fox-72 | 305900 | 253165 | 253165 | 243341 | 238293 |
| Fox-73 | 81854 | 67144 | 67144 | 65553 | 65118 |
| Fox-74 | 41877 | 30720 | 30720 | 29426 | 29414 |
| Fox-75 | 54049 | 46595 | 46595 | 44367 | 44029 |
| Fox-76 | 40592 | 32674 | 32674 | 28896 | 28842 |
| Fox-77 | 53522 | 45815 | 45815 | 44144 | 43847 |
| Fox-78 | 34172 | 20278 | 20278 | 18020 | 18020 |
| Fox-79 | 46726 | 38832 | 38832 | 36464 | 36454 |
| Fox-80 | 40627 | 33650 | 33650 | 31510 | 31510 |
| Fox-81 | 51866 | 44059 | 44059 | 40613 | 40133 |
| Fox-82 | 52180 | 46098 | 46098 | 44370 | 44347 |
| Fox-83 | 42349 | 26558 | 26558 | 24075 | 24026 |
| Fox-84 | 47103 | 43041 | 43041 | 42118 | 42098 |
| Fox-85 | 38988 | 26865 | 26865 | 22961 | 22939 |
| Fox-86 | 45853 | 34332 | 34332 | 31529 | 31446 |
| Fox-87 | 32844 | 22554 | 22554 | 19011 | 18981 |
| Fox-88 | 42103 | 35900 | 35900 | 34356 | 34126 |
| Fox-89 | 59599 | 47730 | 47730 | 45604 | 45536 |
| Fox-90 | 45110 | 34008 | 34008 | 30483 | 30443 |
| Fox-91 | 34284 | 31256 | 31256 | 31015 | 30998 |
| Fox-92 | 54951 | 50373 | 50373 | 49372 | 49230 |
| Fox-93 | 23916 | 20459 | 20459 | 19502 | 19467 |
| Fox-94 | 44742 | 37034 | 37034 | 34514 | 34458 |
| Fox-95 | 56528 | 47656 | 47656 | 45344 | 45132 |
| Fox-96 | 61106 | 37863 | 37863 | 32305 | 32221 |
| Fox-97 | 37973 | 25412 | 25412 | 22631 | 22611 |
| Fox-98 | 40954 | 34261 | 34261 | 32382 | 31814 |
| Fox-99 | 36277 | 23057 | 23057 | 21006 | 20972 |
| Fox-100 | 47965 | 35945 | 35945 | 31170 | 31088 |
| Fox-101 | 45068 | 34508 | 34508 | 32925 | 32832 |
| Fox-102 | 36552 | 27442 | 27442 | 23755 | 23706 |
| Fox-103 | 33182 | 20537 | 20537 | 18753 | 18753 |
| Fox-104 | 54974 | 42379 | 42379 | 39448 | 39050 |
| Fox-105 | 34453 | 28700 | 28700 | 26641 | 26636 |
| Fox-106 | 39177 | 32794 | 32794 | 31045 | 30850 |
| Fox-107 | 34882 | 23259 | 23259 | 21533 | 21520 |
| Fox-108 | 36995 | 34941 | 34941 | 34535 | 34510 |
| Fox-109 | 54753 | 43006 | 43006 | 40689 | 40282 |
| Fox-110 | 68694 | 45353 | 45353 | 39870 | 39761 |
| Fox-111 | 41053 | 26680 | 26680 | 23437 | 23364 |
| Fox-112 | 53157 | 37202 | 37202 | 35855 | 35559 |
| Fox-113 | 50816 | 36473 | 36473 | 33794 | 33754 |
| Fox-114 | 46825 | 35685 | 35685 | 32053 | 30145 |
| Fox-115 | 47090 | 39225 | 39225 | 36200 | 36163 |
| Fox-116 | 34723 | 31011 | 31011 | 30395 | 30356 |
| Fox-117 | 45542 | 35985 | 35985 | 33067 | 32993 |
| Fox-118 | 58638 | 50219 | 50219 | 44120 | 43846 |
| Fox-119 | 32946 | 24979 | 24979 | 22638 | 22638 |
| Fox-120 | 46541 | 39038 | 39038 | 37546 | 37403 |
| Fox-121 | 37361 | 31252 | 31252 | 28973 | 28971 |
| Fox-122 | 56525 | 48085 | 48085 | 46535 | 46307 |
| Fox-123 | 39977 | 36174 | 36174 | 35402 | 35016 |
| Fox-124 | 37856 | 33400 | 33400 | 31263 | 31173 |
| Fox-125 | 75814 | 63308 | 63308 | 58059 | 57733 |
| Fox-126 | 36255 | 26783 | 26783 | 23114 | 23080 |
| Fox-127 | 35914 | 21669 | 21669 | 19255 | 19249 |
| Fox-128 | 43660 | 40482 | 40482 | 39678 | 39652 |
| Fox-129 | 56077 | 38180 | 38180 | 31784 | 31722 |
| Fox-130 | 59345 | 48277 | 48277 | 43811 | 43753 |
| Fox-131 | 57573 | 35635 | 35635 | 32702 | 32647 |
| Fox-132 | 36821 | 28344 | 28344 | 26223 | 26147 |
| Fox-133 | 54072 | 42688 | 42688 | 41824 | 40786 |
| Fox-134 | 40061 | 32177 | 32177 | 30528 | 30472 |
| Fox-135 | 34018 | 27333 | 27333 | 25324 | 25318 |
| Fox-136 | 42870 | 28256 | 28256 | 26642 | 26488 |
| Fox-137 | 39538 | 28066 | 28066 | 25514 | 25468 |
| Fox-138 | 42040 | 29079 | 29079 | 21637 | 21604 |
| Fox-139 | 40528 | 23765 | 23765 | 20985 | 20983 |
| Fox-140 | 48786 | 38044 | 38044 | 33791 | 33481 |
| Fox-141 | 39927 | 23928 | 23928 | 21962 | 21883 |
| Fox-142 | 48466 | 39832 | 39832 | 37191 | 37156 |
| Fox-143 | 41068 | 30958 | 30958 | 28427 | 28383 |
| Fox-144 | 45823 | 35834 | 35834 | 29574 | 29492 |
| Fox-145 | 35400 | 26308 | 26308 | 23743 | 23719 |
| Fox-146 | 43260 | 32919 | 32919 | 28989 | 28958 |
| Fox-147 | 44653 | 29714 | 29714 | 24707 | 24671 |
| Fox-148 | 46027 | 34729 | 34729 | 31334 | 31309 |
| Fox-149 | 34528 | 27429 | 27429 | 22772 | 22760 |
| Fox-150 | 40465 | 31026 | 31026 | 27095 | 27032 |
| Fox-151 | 51017 | 35213 | 35213 | 29175 | 29089 |
| Fox-152 | 50463 | 40348 | 40348 | 39990 | 39567 |
| Fox-153 | 31252 | 24937 | 24937 | 22578 | 22549 |
| Fox-154 | 34316 | 27969 | 27969 | 25850 | 25808 |
| Fox-155 | 26707 | 18135 | 18135 | 14232 | 14220 |
| Fox-156 | 46562 | 32348 | 32348 | 28666 | 28592 |
| Fox-157 | 28960 | 23179 | 23179 | 20531 | 20512 |
| Fox-158 | 80746 | 54957 | 54957 | 46347 | 46259 |
| Fox-159 | 53483 | 40529 | 40529 | 34542 | 34446 |
| Fox-160 | 42856 | 34352 | 34352 | 29921 | 29853 |
| Fox-161 | 55231 | 43090 | 43090 | 37298 | 37256 |
| Fox-162 | 68196 | 36994 | 36994 | 27144 | 27005 |
| Fox-163 | 54925 | 35543 | 35543 | 30227 | 30183 |
| Fox-164 | 56894 | 39366 | 39366 | 33050 | 32871 |
| Fox-165 | 25343 | 17871 | 17871 | 15802 | 15768 |
| Fox-166 | 51886 | 39983 | 39983 | 32522 | 32506 |
| Fox-167 | 40749 | 24917 | 24917 | 22943 | 22924 |
| Fox-168 | 48708 | 47324 | 47324 | 47139 | 47139 |
| Fox-169 | 43353 | 38061 | 38061 | 36394 | 36247 |
| Fox-170 | 44967 | 30234 | 30234 | 27384 | 27293 |
| Fox-171 | 34254 | 28819 | 28819 | 27534 | 27475 |
| Fox-172 | 71879 | 62743 | 62743 | 60599 | 59357 |
| Fox-173 | 57002 | 53711 | 53711 | 52924 | 52468 |
| Fox-174 | 30902 | 22695 | 22695 | 20389 | 20329 |
| Fox-175 | 21820 | 7856 | 7856 | 6413 | 6381 |
| Fox-176 | 46759 | 39459 | 39459 | 37942 | 37602 |
| Fox-177 | 62915 | 26052 | 26052 | 24392 | 24070 |
| Fox-178 | 30033 | 25553 | 25553 | 24031 | 24030 |
| Fox-179 | 81013 | 57627 | 57627 | 54297 | 54167 |
| Fox-180 | 47941 | 24706 | 24706 | 21781 | 21708 |
| Fox-181 | 33711 | 15505 | 15505 | 13917 | 13554 |
| Fox-182 | 55172 | 45133 | 45133 | 41409 | 41269 |
| Fox-183 | 20193 | 16155 | 16155 | 15086 | 15036 |
| Fox-184 | 74180 | 57304 | 57304 | 54711 | 54489 |
| Fox-185 | 62381 | 47631 | 47631 | 40467 | 40371 |
| Fox-186 | 34349 | 29047 | 29047 | 24010 | 23959 |
| Fox-187 | 25742 | 22547 | 22547 | 20316 | 20308 |
| Fox-188 | 47706 | 40799 | 40799 | 38430 | 38395 |
| Fox-189 | 41030 | 31001 | 31001 | 28131 | 28108 |
| Fox-190 | 31460 | 10704 | 10704 | 9247 | 9240 |
| Fox-191 | 25173 | 20006 | 20006 | 17183 | 17169 |
| Fox-192 | 29451 | 20726 | 20726 | 18668 | 18579 |
| Fox-193 | 30277 | 22979 | 22979 | 21257 | 21224 |
| Fox-194 | 40723 | 31060 | 31060 | 28743 | 28701 |
| Fox-195 | 30398 | 23700 | 23700 | 21579 | 21574 |
| Fox-196 | 61113 | 47340 | 47340 | 42980 | 42526 |
| Fox-197 | 34188 | 29549 | 29549 | 26701 | 26607 |
| Fox-198 | 49929 | 36417 | 36417 | 33505 | 33458 |
| Fox-199 | 34041 | 29146 | 29146 | 22259 | 22233 |
| Fox-200 | 13276 | 9419 | 9419 | 7756 | 7748 |
| Fox-201 | 43830 | 39024 | 39024 | 38141 | 38096 |
| Fox-202 | 19195 | 9881 | 9881 | 8961 | 8945 |
| Fox-203 | 18022 | 16665 | 16665 | 16079 | 16079 |
| Fox-204 | 28353 | 24252 | 24252 | 21643 | 21639 |
| Fox-205 | 34793 | 30080 | 30080 | 27963 | 27840 |
| Fox-206 | 49186 | 33327 | 33327 | 29129 | 28981 |
| Fox-207 | 35453 | 30980 | 30980 | 29246 | 28861 |
| Fox-208 | 17955 | 13526 | 13526 | 12293 | 12289 |
| Fox-209 | 31693 | 20183 | 20183 | 18463 | 18420 |
| Fox-210 | 53426 | 42493 | 42493 | 38621 | 38154 |
| Fox-211 | 29512 | 22583 | 22583 | 19457 | 19390 |
| Fox-212 | 51686 | 47088 | 47088 | 46614 | 45695 |
| Fox-213 | 20055 | 18481 | 18481 | 17379 | 17337 |
| Fox-214 | 72433 | 69465 | 69465 | 68802 | 66775 |
| Fox-215 | 59497 | 40724 | 40724 | 39806 | 38914 |
| Fox-216 | 26999 | 22453 | 22453 | 20133 | 20098 |
| Fox-217 | 47723 | 41067 | 41067 | 40265 | 39625 |
| Fox-218 | 29212 | 27657 | 27657 | 26997 | 26774 |
| Fox-219 | 62736 | 50784 | 50784 | 49737 | 48587 |
| Fox-220 | 43094 | 27401 | 27401 | 26902 | 26782 |
| Fox-221 | 40670 | 34942 | 34942 | 28693 | 28670 |
| Fox-222 | 19018 | 16703 | 16703 | 15692 | 15682 |
| Fox-223 | 28146 | 25865 | 25865 | 25070 | 25055 |
| Fox-224 | 17646 | 15756 | 15756 | 15155 | 15146 |
| Fox-225 | 45348 | 31986 | 31986 | 29195 | 29153 |
| Fox-226 | 38755 | 31745 | 31745 | 30051 | 29989 |
| Fox-227 | 36484 | 35061 | 35061 | 34298 | 33480 |
| Fox-228 | 45449 | 43897 | 43897 | 43397 | 43312 |
| Fox-229 | 25604 | 23181 | 23181 | 22448 | 22417 |
| Fox-230 | 39074 | 32852 | 32852 | 31028 | 30986 |
| Fox-231 | 43365 | 41348 | 41348 | 40778 | 39758 |
| Fox-232 | 37706 | 23621 | 23621 | 20594 | 20551 |
| Fox-233 | 21546 | 19495 | 19495 | 18453 | 18430 |
| Fox-234 | 44843 | 42915 | 42915 | 42398 | 42250 |
| Fox-235 | 43203 | 41909 | 41909 | 41514 | 41062 |
| Fox-236 | 36824 | 28420 | 28420 | 25209 | 25199 |
| Fox-237 | 60625 | 58166 | 58166 | 57267 | 55719 |
| Fox-238 | 52206 | 50202 | 50202 | 49530 | 48562 |
| Fox-239 | 52511 | 43538 | 43538 | 42318 | 42085 |
| Fox-240 | 16308 | 12420 | 12420 | 12076 | 12076 |
| Fox-241 | 21347 | 15934 | 15934 | 14554 | 14551 |
| Fox-242 | 37563 | 34442 | 34442 | 26748 | 26528 |
| Fox-243 | 32535 | 23399 | 23399 | 21042 | 21002 |
| Fox-244 | 51084 | 37316 | 37316 | 33674 | 33648 |
| Fox-245 | 57302 | 42056 | 42056 | 38826 | 38792 |
| Fox-246 | 35646 | 24203 | 24203 | 22297 | 22260 |
| Fox-247 | 38441 | 28161 | 28161 | 26273 | 26207 |
| Fox-248 | 50351 | 48085 | 48085 | 47436 | 46818 |
| Fox-249 | 37599 | 31380 | 31380 | 28709 | 28635 |
| Fox-250 | 32592 | 30117 | 30117 | 29340 | 28979 |
| Fox-251 | 18205 | 16955 | 16955 | 16459 | 16378 |
| Fox-252 | 14547 | 11939 | 11939 | 11570 | 11567 |
| TOTAL | 11493554 | 9055526 | 9055526 | 8406012 | 8354655 |
| AVERAGE | 45250.21 | 35651.68 | 35651.68 | 33094.54 | 32892.34 |
| † *These samples were sequenced twice* | | |  |  |  |

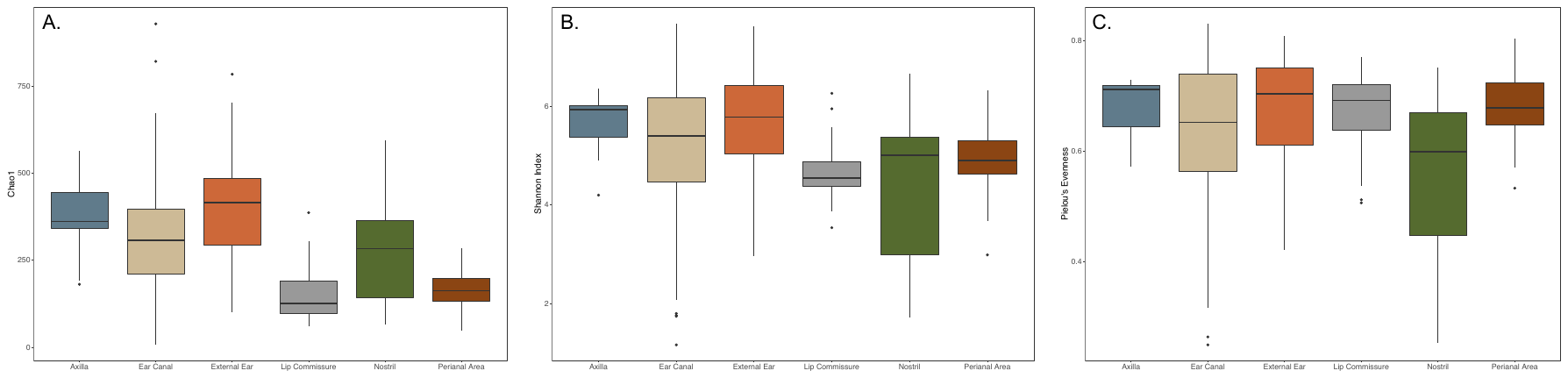

**Supplemental Figure S2.** Boxplots of three alpha diversity metrics rarefied to 6381: (A) Chao1 Index, (B) Shannon Index, and (C) Pielou’s Evenness Metric. Alpha diversity was significantly different between body sites using the Chao 1 (Kruskal-Wallis test; *H*=69.852, *p*<0.001) and Shannon (*H*= 23.242, *p*<0.001) indices, whereas Pielou’s Evenness did not differ significantly between sites (*H*=8.800, *p*=0.117).

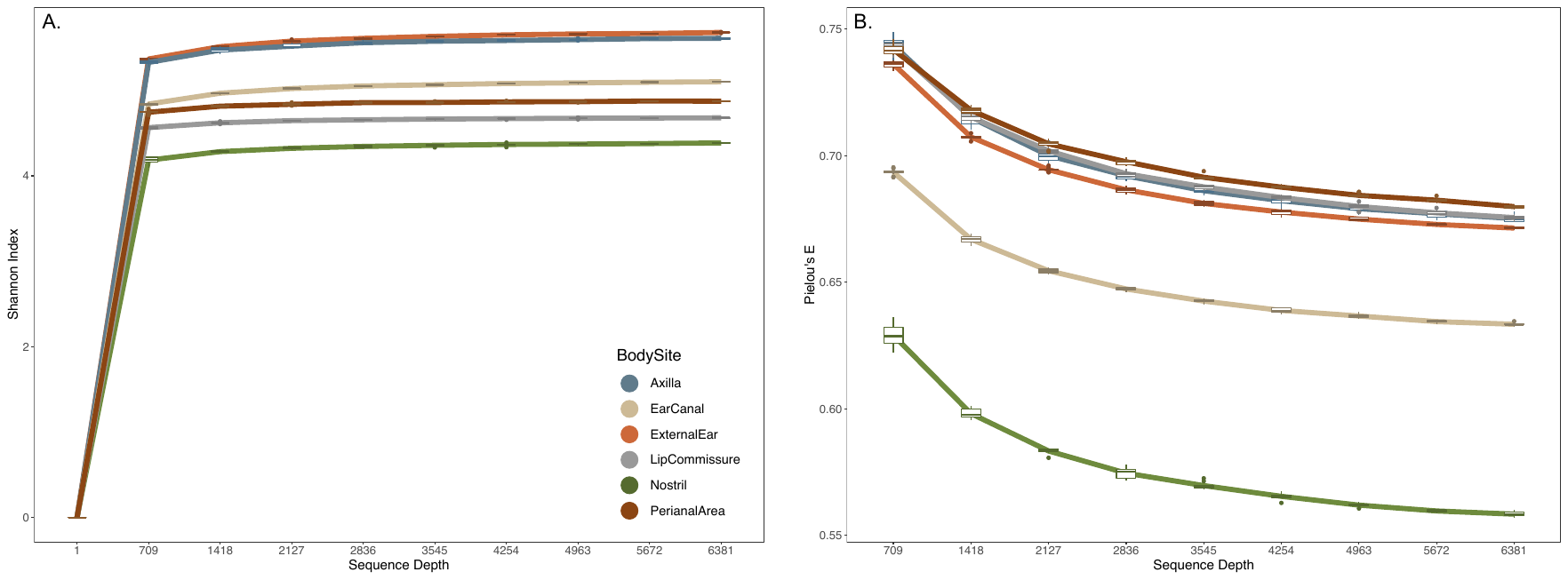

**Supplemental Figure S3.** Rarefaction curves of two alpha diversity metrics: (A) Shannon Index and (B) Pielou’s Evenness Metric. Lines represent mean values across samples and iterations (n=10), with box plots representing mean values across samples for each iteration.

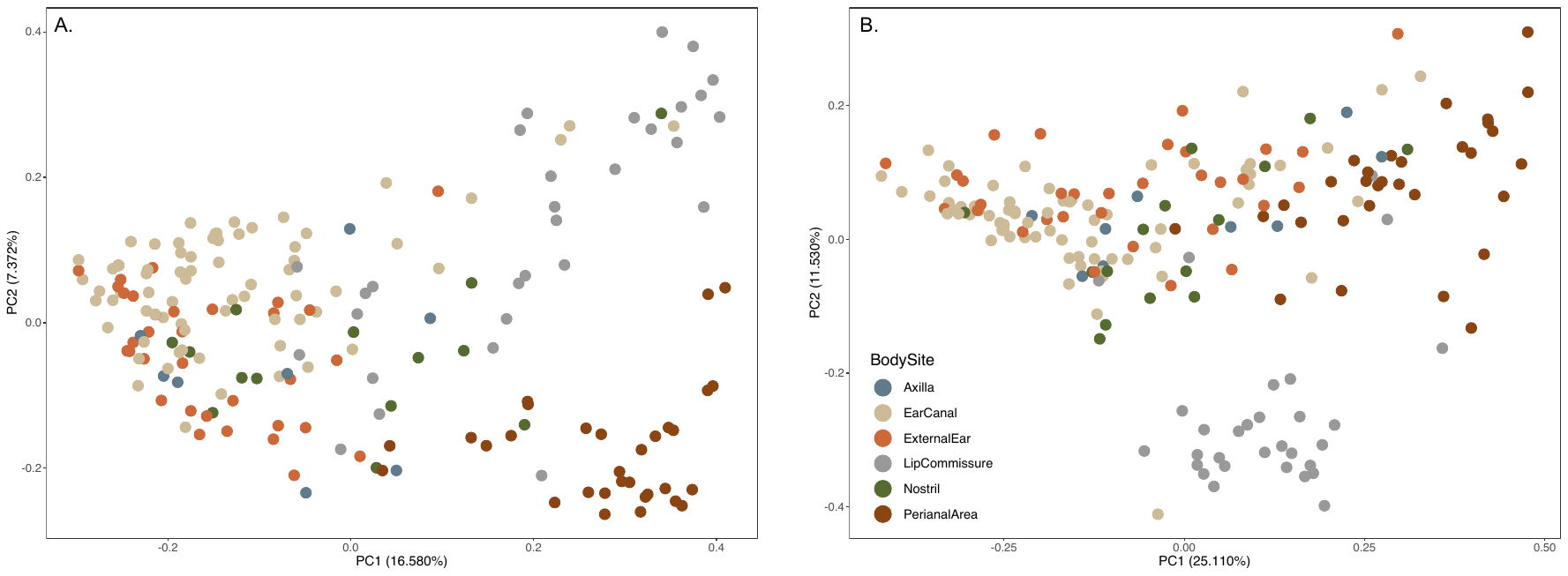

**Supplemental Figure S4.** Principal coordinate analysis (PCoA) plots using two phylogeny-based distance measures: (A) unweighted and (B) weighted UniFrac distances. Beta diversity significantly differed between body sites using both unweighted (*PERMANOVA; pseudo-F*=7.990, *p*=0.001) and weighted (*pseudo-F*=12.474, *p*=0.001) distances.

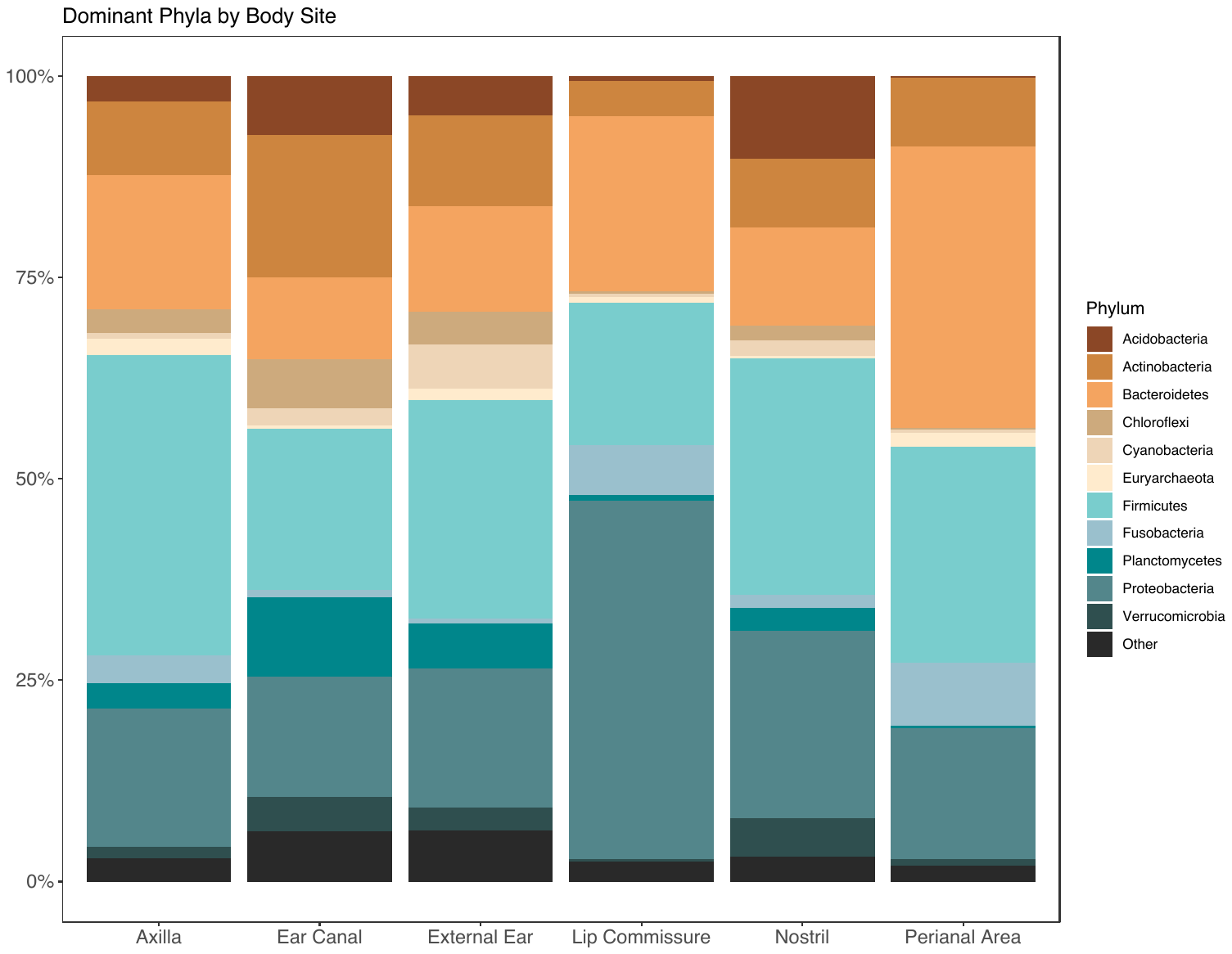

**Supplemental Figure S5.** Taxonomic composition at each body site at the phylum level.

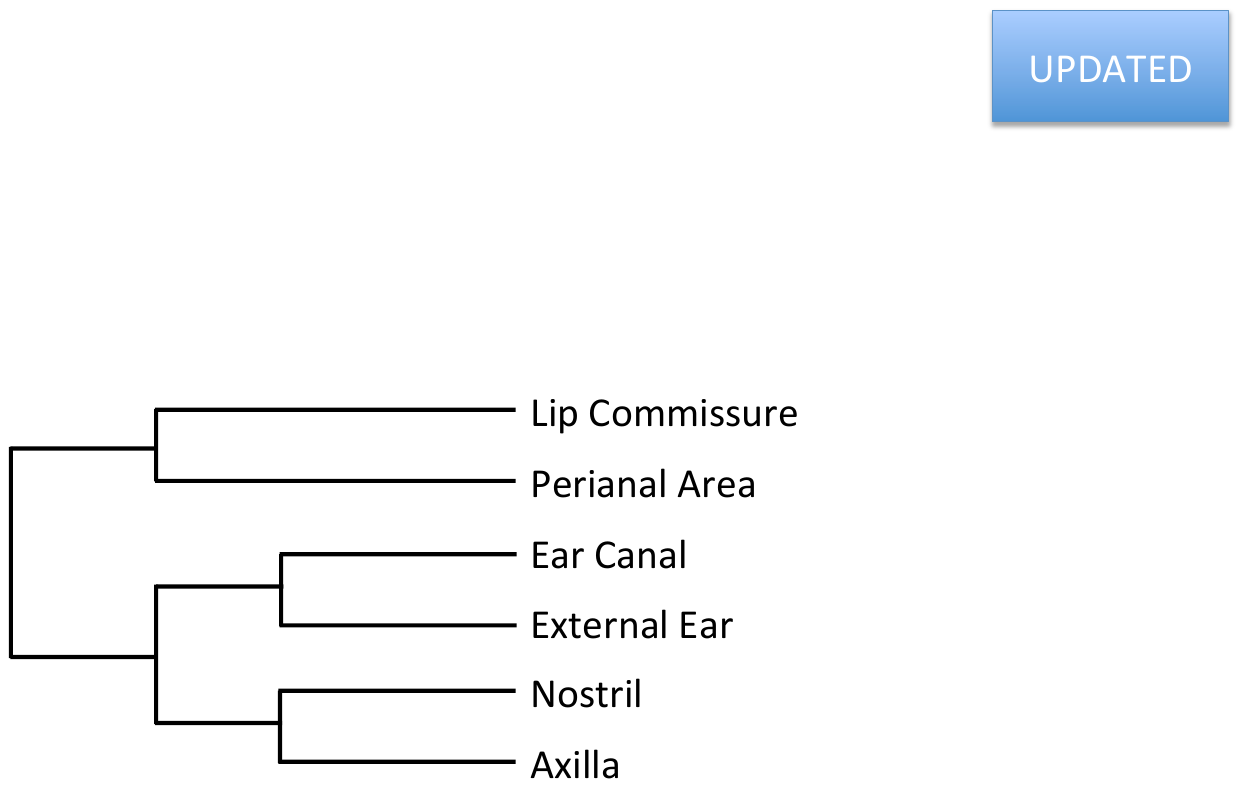

**Supplemental Figure S6.** Hierarchical clustering of body sites based on Bray-Curtis and Euclidean distance metrics and the “average” linkage method.

**Supplemental Table S2.** Results from Chi-square tests of independence for every pairwise combination of ear canal phenotypes. Sample size for each presence class is given in parentheses. Significance was determined via modified-FDR correction (corrected *p*=0.015). With the exception of nodule presence (where sample size was heavily skewed towards the absence class), all phenotypes were significantly correlated.

| **Phenotype (present, absent)** | **Wax (58, 60)** | ***Liquor Puris* (19, 99)** | **Nodules (8, 110)** | **Pigment (45, 73)** | **Inflammation (44, 74)** |
| --- | --- | --- | --- | --- | --- |
| **Mites (42, 76)** | *X^2^*=58.316, *p<*0.001* | *X^2^*=9.008, *p*=0.003* | *X^2^*=1.597, *p*=0.206 | *X^2^*=47.894, *p<*0.001* | *X^2^*=22.158, *p*<0.001* |
| **Wax (58, 60)** | . | *X^2^*=12.871, *p<*0.001* | *X^2^*=1.319, *p*=0.251 | *X^2^*=48.561, *p*<0.001* | *X^2^*=32.075, *p<*0.001* |
| ***Liquor Puris* (19, 99)** | . | . | *X^2^*=4.856, *p*=0.028 | *X^2^*=13.993, *p*<0.001* | *X^2^*=7.867, *p*=0.005* |
| **Nodules (8, 110)** | . | . | . | *X^2^*=6.762, *p*=0.009* | *X^2^*=7.093, *p*=0.008* |
| **Pigment (45, 73)** | . | . | . | . | *X^2^*=33.286, *p*<0.001* |
| ** p-value below the FDR corrected threshold of 0.015; N.B., one sample was removed from these analyses due to missing phenotypes* | | | | |  |

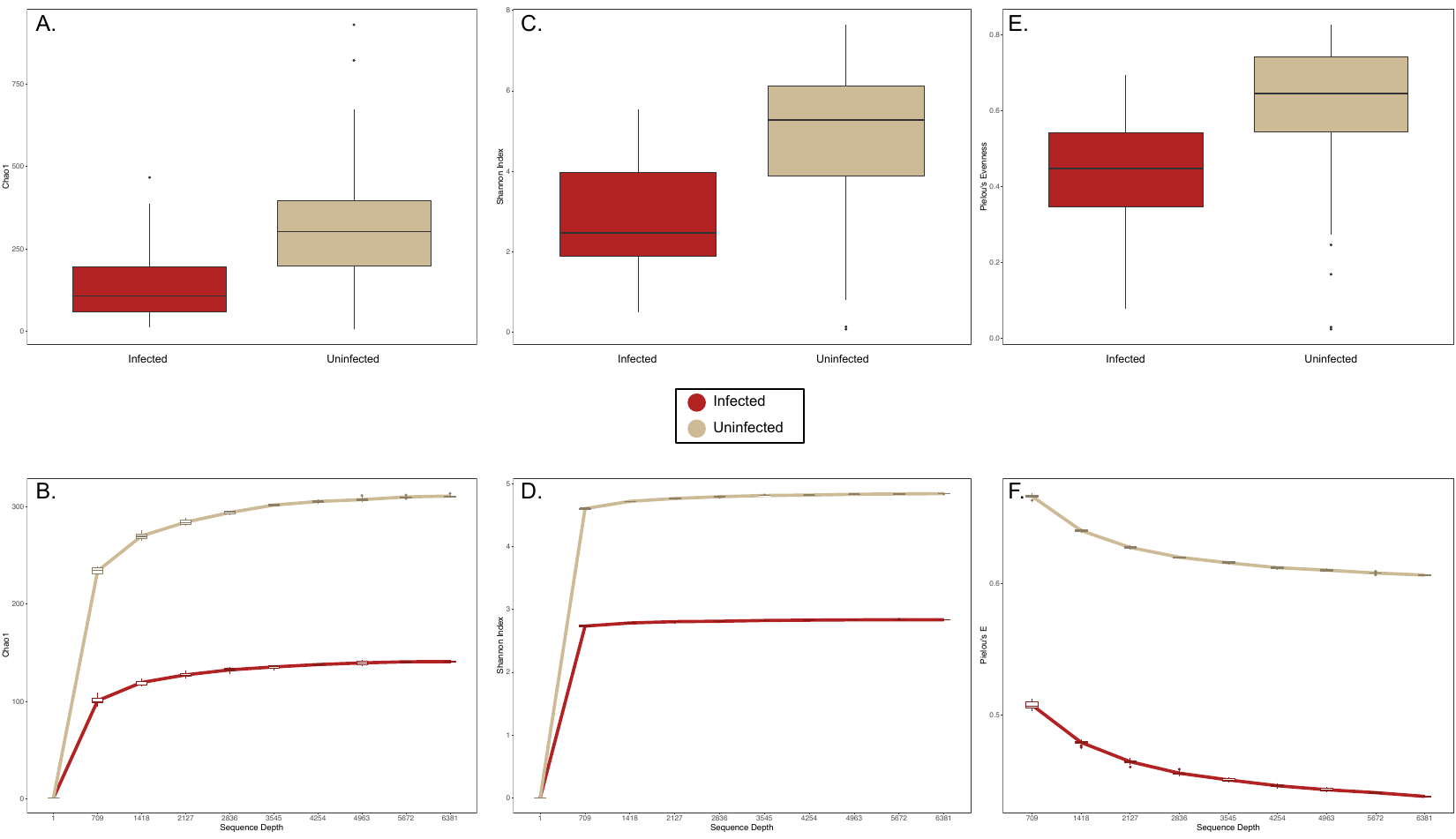

**Supplemental Figure S7.** Paired boxplots and rarefaction curves of three alpha diversity metrics: (A-B) Chao1 Index, (C-D) Shannon Index, and (E-F) Pielou’s Evenness Metric. Mite-infected ear canals exhibited significantly reduced diversity using all three metrics (Kruskal-Wallis Test; Chao1, *H*=25.635, *p*<0.001; Shannon Index, *H*=31.481, *p*<0.001; Pielou’s Evenness Metric, *H*=27.848, *p*<0.001). In rarefaction plots, lines represent mean values across samples and iterations (n=10), with box plots representing mean values across samples for each iteration.

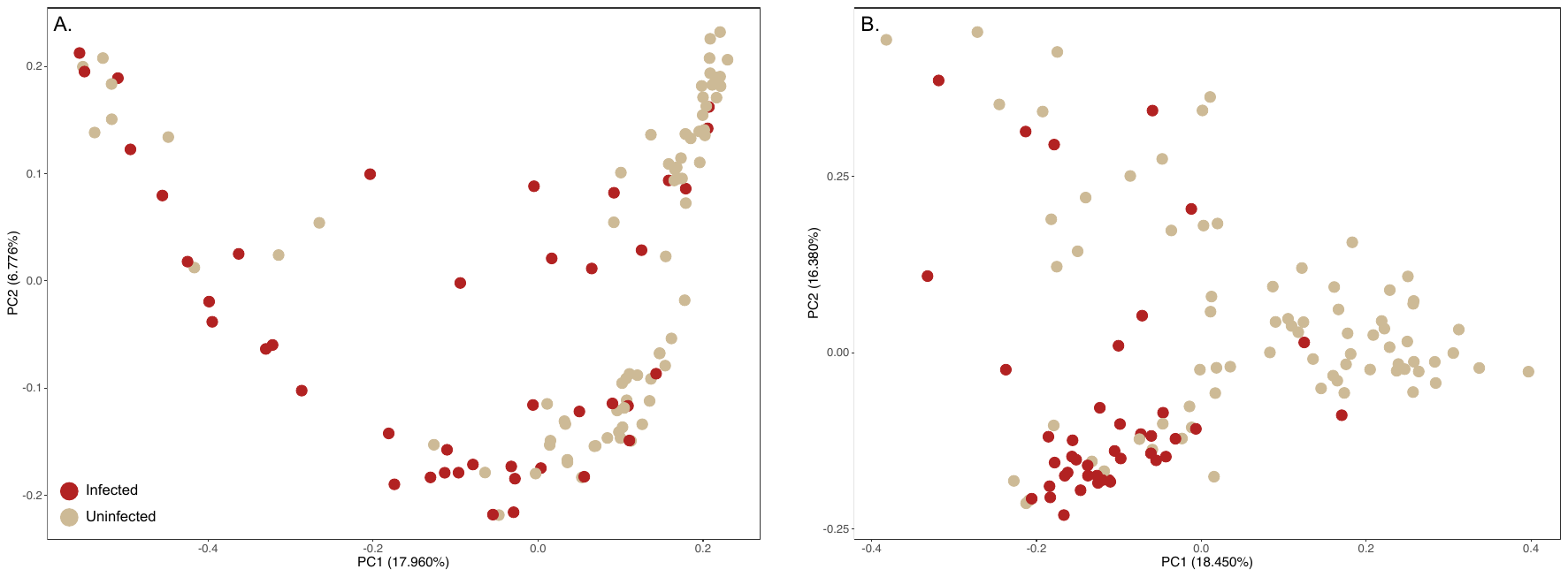

**Supplemental Figure S8.** Principal coordinate analysis (PCoA) plots using two phylogeny-based distance measures: (A) unweighted and (B) weighted UniFrac distances. Beta diversity significantly differed between body sites using both unweighted (*PERMANOVA; pseudo-F*=4.281, *p*=0.001) and weighted (*pseudo-F*=8.942, *p*=0.001) distances.

**Supplemental Table S3.** Top 25 NCBI BLAST results for the feature identified through ANCOM as differentially abundant between mite-infected and uninfected ear canals.

| **Species** | **Max Score** | **E value** | **% Identity** | **Accession No.** |
| --- | --- | --- | --- | --- |
| *Staphylococcus pseudintermedius* | 410 | 9.00E-111 | 100% | CP035740.1 |
| *Staphylococcus pseudintermedius* | 410 | 9.00E-111 | 100% | CP035741.1 |
| *Staphylococcus pseudintermedius* | 410 | 9.00E-111 | 100% | CP035742.1 |
| *Staphylococcus pseudintermedius* | 410 | 9.00E-111 | 100% | CP035743.1 |
| *Staphylococcus pseudintermedius* | 410 | 9.00E-111 | 100% | CP032682.1 |
| *Uncultured Staphylococcus sp.* | 410 | 9.00E-111 | 100% | MH728109.1 |
| *Staphylococcus pseudintermedius* | 410 | 9.00E-111 | 100% | MK447575.1 |
| *Staphylococcus delphini* | 410 | 9.00E-111 | 100% | MK396462.1 |
| *Staphylococcus schleiferi* | 410 | 9.00E-111 | 100% | CP035007.1 |
| *Staphylococcus pseudintermedius* | 410 | 9.00E-111 | 100% | AP019372.1 |
| *Staphylococcus aureus* | 410 | 9.00E-111 | 100% | LR134267.1 |
| *Staphylococcus hyicus* | 410 | 9.00E-111 | 100% | LR134264.1 |
| *Staphylococcus delphini* | 410 | 9.00E-111 | 100% | LR134263.1 |
| *Staphylococcus pseudintermedius* | 410 | 9.00E-111 | 100% | MH392298.1 |
| *Staphylococcus pseudintermedius* | 410 | 9.00E-111 | 100% | MF185092.1 |
| *Staphylococcus muscae* | 410 | 9.00E-111 | 100% | CP027848.1 |
| *Staphylococcus felis* | 410 | 9.00E-111 | 100% | CP027770.1 |
| *Staphylococcus pseudintermedius* | 410 | 9.00E-111 | 100% | MG576276.1 |
| *Staphylococcus pseudintermedius* | 410 | 9.00E-111 | 100% | MG576275.1 |
| *Staphylococcus pseudintermedius* | 410 | 9.00E-111 | 100% | MG576274.1 |
| *Staphylococcus muscae* | 410 | 9.00E-111 | 100% | LT906464.1 |
| *Staphylococcus lutrae* | 410 | 9.00E-111 | 100% | CP020773.1 |
| *Uncultured Staphylococcus sp.* | 410 | 9.00E-111 | 100% | KX833847.1 |
| *Staphylococcus pseudintermedius* | 410 | 9.00E-111 | 100% | CP015626.1 |
| *Staphylococcus pseudintermedius* | 410 | 9.00E-111 | 100% | CP016073.1 |

**Supplemental Table S4.** Percent difference in *r^2^* calculated between the full OLS model and one excluding a single variable at a time. Mite infection is the variable of largest effect in this model, since its removal caused the largest change in *r^2^*.

| **Variable** | **% difference in *r^2^*** |
| --- | --- |
| Sampling Location | 1.40% |
| Age Class | 1.47% |
| Body Condition | 1.09% |
| Sex | 1.11% |
| Weight | 1.45% |
| Mite Infection | 2.28% |
