## Supporting Information for "Microbial dysbiosis and its implications for disease in a genetically depauperate species"

**Supporting Information 2: Subsampled Ear Canal Analyses**

**Introduction**

The full ear canal dataset consisted of 119 samples collected from 73 foxes. Since each individual was sampled 1-3 times, we randomly subsampled our dataset (73 samples total, 1 sample per fox) and re-ran all analyses to confirm the robustness of our results. Please refer to the **Materials and Methods** described in the manuscript for details on how each test was run.

**Results**

(1) Alpha diversity metrics showed the same pattern of decreased species richness (Kruskall-Wallis Test; Chao 1 Index, *H*=13.882, *p*<0.001), species abundance and evenness (Shannon, *H*=17.030, *p*<0.001), and species equitability (Pielou’s E, *H*=13.494, *p*<0.001) in mite-infected ear canals (Supplemental Figure SE1).

(2) Beta diversity metrics measuring species presence and abundance also differed between the two infection groups. Though the clustering was less clear in the PCoA plots (Supplemental Figure SE2), *PERMANOVA* results were significant when using the (A) Bray-Curtis (*pseudo-F*= 4.659 *p*=0.001), (B) unweighted UniFrac (*pseudo-F*=2.443, *p*=0.003), and (C) weighted UniFrac (*pseudo-F*=4.894, *p*=0.001) distance measures.

(3) The taxonomic composition of mite infected and uninfected ear canals also matched the results obtained with the full dataset, with Bacilli showing a much higher relative abundance in mite infected ear canals (40.05%) compared to uninfected ear canals (16.89%; Supplemental Figure SE3).

(4) To formally test this relationship, we ran ANCOM and gneiss balances on our subsampled feature table, which we further filtered for a minimum frequency of 50 and a minimum number of samples of 10.

ANCOM again returned a single feature as significantly differentially abundant between infection groups: 3f0449c545626dd14b585e9c7b2d16f4 (*W*=210). This was the same feature identified using our full dataset, and matched *Staphylococcus pseudintermedius* with 100% identity when we input the sequence into NCBI’s BLASTn tool.

We then ran an ordinary least squares (OLS) regression on the gneiss balances calculated with the subsampled dataset. As with the full dataset, we set sampling area, sex, age class, weight, body condition, and presence of ear mites as fixed effects. The overall model explained 11.32% of the variation, with ear mite infection the variable of largest effect (2.81%; Supplemental Table SE1). Balance y02 was statistically significant with respect to ear mite infection (*p*<0.001). Visualizing the taxonomic make up of this balance showed that Bacilli was the only class of microbes with drastically different proportions between infection groups (Supplemental Figure SE4). The feature underlying this pattern was 3f0449c545626dd14b585e9c7b2d16f4, previously identified as *S. pseudintermedius*. In this subsampled analysis, correlation-clustering grouped this feature with one other in the tree hierarchy: 13e52764e1c08c4fd9902769bf9e022e, which matched to *Enterococcus spp.* using NCBI BLASTn (Altschul et al. 1990). The result that *S. pseudintermedius* was the primary taxon driving observed changes is therefore consistent between ANCOM and gneiss using the full and subsampled datasets.

One difference concerns the additional taxa grouped with *S. pseudintermedius* in the tree hierarchy. The full dataset returned *Enterococcus spp.* (Bacilli) and *Corynebacterium spp*. (Actinobacteria), whereas the subsampled dataset only returned *Enterococcus spp.* This suggests that the higher proportion of Anctinobacteria observed with the full dataset was perhaps driven by (1) specific foxes sampled multiple times or (2) specific samples that were excluded from the subsampled analysis by random chance.

**Discussion**

Results obtained with our full (n=119) and subsampled (n=73) datasets were nearly identical. Alpha diversity was significantly lower in mite-infected ear canals, community composition was significantly different between infection groups, and Bacilli (more specifically, *S. pseudintermedius*) had a significantly higher relative abundance in mite-infected ear canals when compared to uninfected ear canals. Regression models returned only one balance with significant differences with respect to ear mite infection, and the same conclusion was drawn from each: *S. pseudintermedius* had higher relative abundance in mite-infected ear canals. As almost all subsampled results matched the full dataset, we retained all samples for analyses presented in the main manuscript.

**Tables and Figures**

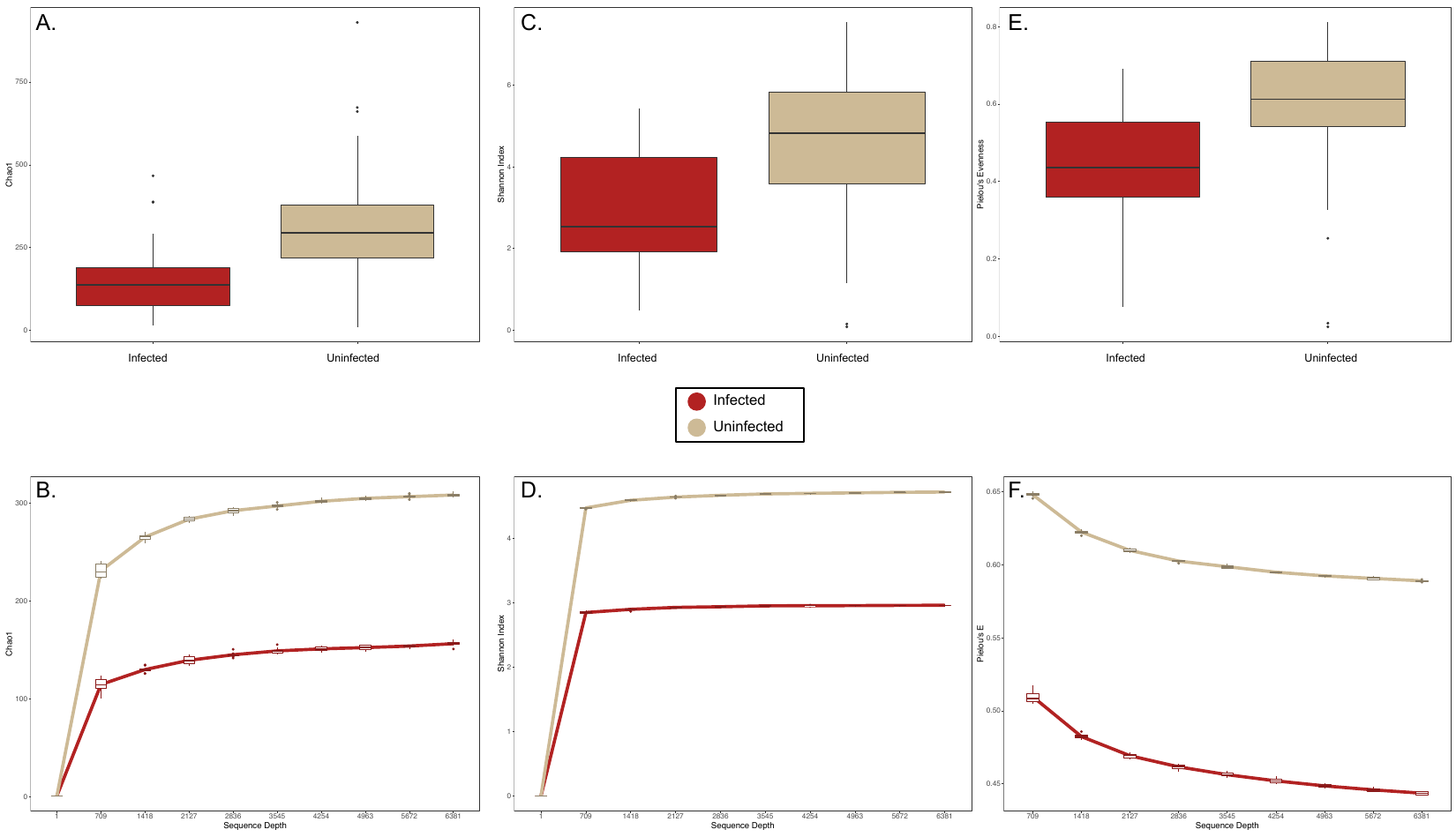

**Supplemental Figure SE1.** Paired boxplots and rarefaction curves of three alpha diversity metrics: (A-B) Chao1 Index, (C-D) Shannon Index, and (E-F) Pielou’s Evenness Metric. Mite-infected ear canals exhibited significantly reduced diversity using all three metrics (Kruskall-Wallis Test; Chao1, *H*=13.882, *p*<0.001; Shannon Index, *H*=17.030, *p*<0.001; Pielou’s Evennesss Metric, *H*=13.494, *p*<0.001). In rarefaction plots, lines represent mean values across samples and iterations (n=10), with box plots representing mean values across samples for each iteration.

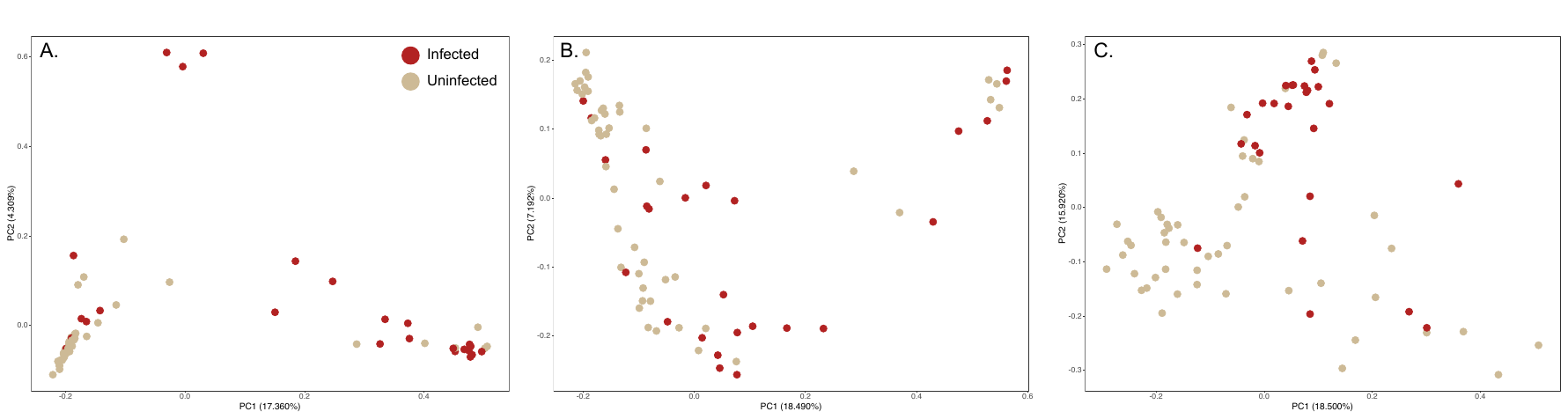

**Supplemental Figure SE2.** Principal coordinate analysis plots using three distance measures: (A) Bray-Curtis dissimilarity index and phylogeny-based (B) unweighted and (B) weighted UniFrac distances. Significant differences were observed using Bray-Curtis (*PERMANOVA; pseudo-F*= 4.659 *p*=0.001), (B) unweighted UniFrac (*pseudo-F*=2.443, *p*=0.003), and (C) weighted UniFrac (*pseudo-F*=4.894, *p*=0.001) distance measures.

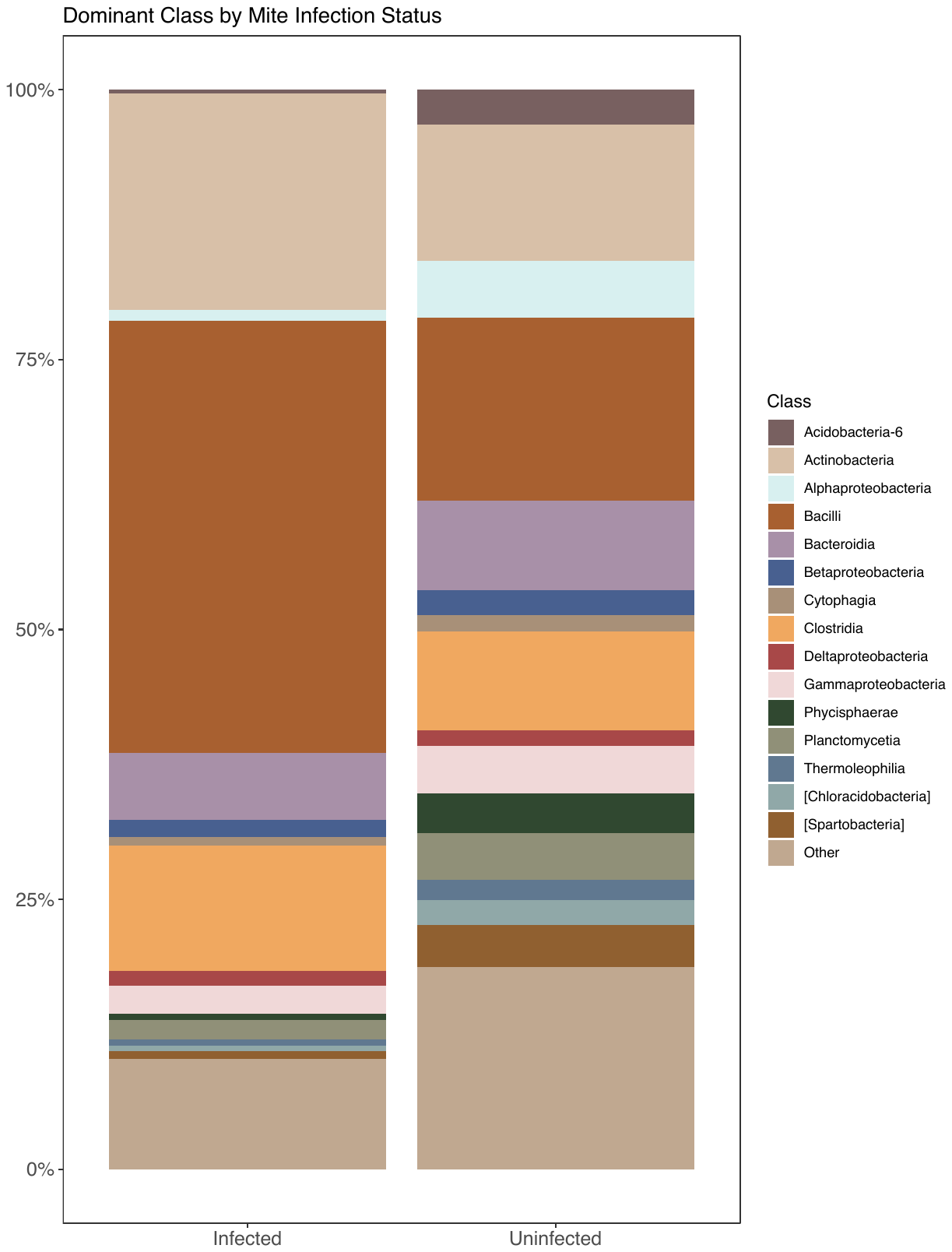

**Supplemental Figure SE3.** Taxonomic composition exhibited the same classes of microbes in both infection groups, with Bacilli far more abundant in mite-infected ear canals.

**Supplemental Table SE1**. Percent difference in *r^2^* calculated between the full OLS model and one excluding a single variable at a time. Mite infection was the variable of largest effect in this model, since its removal caused the largest change in *r^2^*.

| **Variable** | **% difference in *r^2^*** |
| --- | --- |
| Sampling Location | 1.88% |
| Age Class | 1.62% |
| Body Condition | 1.31% |
| Sex | 1.31% |
| Weight | 1.95% |
| Mite Infection | 2.81% |

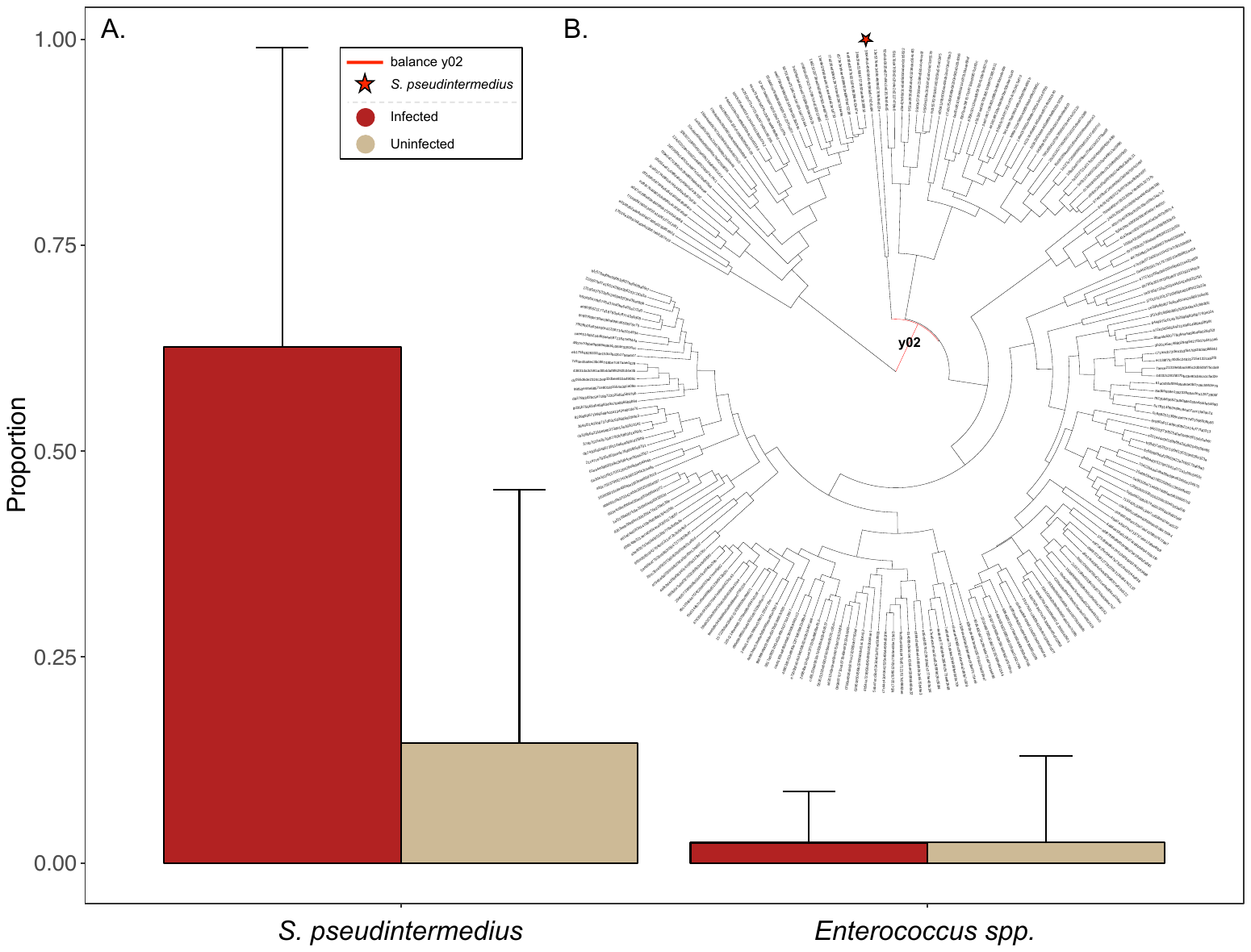

**Supplemental Figure SE4.** (A) Proportion bar plots for two taxonomic features contained in the gneiss balance associated with ear mite infection: balance y02. *S. pseudintermedius* had the largest difference in proportion between mite-infected and uninfected ear canals, with *Enterococcus spp.* clustered with *S. pseudintermedius* in the (B) hierarchy relating all microbial taxa. Balances y02 is highlighted in red, and *S. pseudintermedius* is indicated with a star.
